# Role of BMP signaling during early development of the annelid *Capitella teleta*

**DOI:** 10.1101/2020.11.15.383695

**Authors:** Nicole B. Webster, Michele Corbet, Abhinav Sur, Néva P. Meyer

**Author notes:** corresponding author, Clark University, Biology Department, 950 Main Street, Worcester, MA 01610-1400.

## Abstract

The mechanisms regulating nervous system development are still unknown for a wide variety of taxa. In insects and vertebrates, bone morphogenetic protein (BMP) signaling plays a key role in establishing the dorsal-ventral (D-V) axis and limiting the neuroectoderm to one side of that axis, leading to speculation about the conserved evolution of centralized nervous systems. Studies outside of insects and vertebrates show a more diverse picture of what, if any role, BMP signaling plays in neural development across Bilateria. This is especially true in the morphologically diverse Spiralia (~Lophotrochozoa). Despite several studies of D-V axis formation and neural induction in spiralians, there is no consensus for how these two processes are related, or whether BMP signaling may have played an ancestral role in either process. To determine the function of BMP signaling during early development of the spiralian annelid *Capitella teleta*, we incubated embryos and larvae in BMP4 protein for different amounts of time. Adding exogenous BMP protein to early-cleaving *C. teleta* embryos had a striking effect on formation of the brain, eyes, foregut, and ventral midline in a time-dependent manner. However, adding BMP did not block brain or VNC formation or majorly disrupt the D-V axis. We identified three key time windows of BMP activity. 1) BMP treatment around birth of the 3^rd^-quartet micromeres caused the loss of the eyes, radialization of the brain, and a reduction of the foregut, which we interpret as a loss of A- and C-quadrant identities with a possible trans-fate switch to a D-quadrant identity. 2) Treatment after birth of micromere 4d induced formation of a third ectopic brain lobe, eye, and foregut lobe, which we interpret as a trans-fate switch of B-quadrant micromeres to a C-quadrant identity. 3) Continuous BMP treatment from late cleavage (4d + 12h) through mid-larval stages resulted in a modest expansion of *Ct-chrdl* expression in the dorsal ectoderm and a concomitant loss of the ventral midline (neurotroch ciliary band). Loss of the ventral midline was accompanied by a collapse of the bilaterally-symmetric ventral nerve cord, although the total amount of neural tissue did not appear to be greatly affected. Our results compared to those from other annelids and molluscs suggest that BMP signaling was not ancestrally involved in delimiting neural tissue to one region of the D-V axis. However, the effects of ectopic BMP on quadrant-identity during cleavage stages may represent a non-axial organizing signal that was present in the last common ancestor of annelids and mollusks. Furthermore, in the last common ancestor of annelids, BMP signaling may have functioned in patterning ectodermal fates along the D-V axis in the trunk. Ultimately, studies on a wider range of spiralian taxa are needed to determine the role of BMP signaling during neural induction and neural patterning in the last common ancestor of this group. Ultimately, these comparisons will give us insight into the evolutionary origins of centralized nervous systems and body plans.

## Introduction

Nervous systems are a key feature of most animal taxa, but how evolution produced the wide variety of extant nervous systems is still unknown. Morphologically, nervous systems range from those that are predominantly a plexus (Richter et al., 2010), such as in ctenophores and cnidarians (Hejnol and Rentzsch, 2015; Schmidt-Rhaesa et al., 2015), to those that have a high degree of centralization, e.g. the ‘centralized nervous systems’ (CNSs) of annelids, insects, and chordates (Arendt et al., 2008; Holland, 2003). The morphological organization of a nervous system can also have functional implications, such as how signals for sensory input and motor output are processed and coordinated. What mechanisms regulate nervous system development, and especially how that differs between taxa, is of particular interest to understand evolution of this complex system.

In insects and vertebrates, the CNS arises from a region of ectoderm that is induced to become neural as part of dorsal-ventral (D-V) axis specification, although the neuroectoderm is dorsal in vertebrates and ventral in insects. D-V axis formation relies in part on a gradient of Bone Morphogenetic Protein (BMP) signaling, with high levels of BMP signaling inhibiting neural induction (De Robertis and Kuroda, 2004; Holley et al., 1995; Wilson and Edlund, 2001). The similarities between vertebrate and insect neural induction, where BMP signaling plays an anti-neural role, have prompted speculation that the last common ancestor of Bilateria had a CNS that was localized along a D-V axis (Arendt and Nubler-Jung 1999; Denes et al. 2007; De Robertis 2008). However, comparison of data from a wider range of taxa from deuterostomes and ecdysozoans shows less support for this hypothesis (see Discussion).

Examining new systems can lead to a new perspective into neural development and evolution. Spiralia (≈Lophotrochozoa) includes taxa with widely divergent body plans such as annelids, molluscs, brachiopods, rotifers, and platyhelminths (Kocot et al., 2016). These diverse morphologies include a great variety of nervous systems, from the highly intelligent brains of cephalopods (Shigeno et al., 2018) to the rope-ladder-like ventral nerve cords in many annelids (Helm et al., 2018) to a single subesophageal ganglion in inarticulate brachiopods (Nielsen, 2005). Such diversity leads to opportunities for comparative studies to understand the evolutionary origins of CNSs. Spiralians have an added benefit for studying the evolution of body plans, including the CNS. Developmentally many taxa share an ancestral stereotyped cleavage program called spiral cleavage where each cell (blastomere) acquires distinct fates during embryogenesis that contributes to a specific set of tissues (Henry, 2014).

Despite several studies of D-V axis formation and neural induction in spiralians, there is no consensus for how these two processes are related, or whether BMP signaling may have played an ancestral role in either process. In the mollusc *Tritia* (previously *Ilyanassa*) *obsoleta*, knocking down BMP signaling during early cleavage stages resulted in a similar phenotype to removing the organizer signal (i.e. removing the polar lobe), including a disrupted D-V axis. In contrast, adding BMP4 protein partially rescued the removal of the polar lobe (Lambert et al., 2016). While BMP signaling appears to be part of the axial organizer in this animal, exogenous BMP protein caused ectopic eye and likely brain formation rather than a loss of neural tissue as seen in vertebrates and insects, and no effect on trunk neural tissue was reported (Lambert et al., 2016). These data suggest that BMPs may actually induce neural tissue in *T. obsoleta* rather than block it as in vertebrates and insects. In *Crepidula fornicata*, another mollusc, BMP protein induced a pinching off of part of the episphere, while the trunk was relatively normal with a D-V axis and neural tissue (Lyons et al., 2020).

Annelids have a CNS comprising an anterior brain and a ventral nerve cord (VNC) (Muller, 2006), but it is unclear if BMP signaling is involved in early D-V axis formation and/or neural specification of the CNS. In the leech *Helobdella*, gain- and loss-of-function experiments demonstrated that BMP signaling is important for D-V patterning within the ectoderm; however, no effect on neural induction was reported (Kuo and Weisblat, 2011). In the annelid *Platynereis dumerilii*, application of BMP4 protein to early trochophores affected D-V patterning of the ectoderm and VNC, but did not repress neural induction (Denes et al., 2007). It is not clear if either study manipulated BMP signaling early enough to affect D-V axis formation (as opposed to localized patterning of the ectoderm) or neural induction. In the annelid *Capitella teleta*, recent studies have shown that Activin/Nodal signaling, not BMP, are responsible for D-V axis formation, but the function in neural specification is still undetermined (Lanza and Seaver, 2018, 2020a). This leaves open the question of whether or not BMP signaling was ancestrally involved in these processes in annelids.

Here we further explore the role of BMP signaling during neural development in the annelid *C. teleta*. By exposing cleavage-stage embryos at different timepoints to exogenous BMP protein, we show the importance of timing in the role BMP plays in neural development, as well as key differences between the effects on the trunk and episphere. BMP treatment at early cleavage stages, just after axial organizer signaling, resulted in animals with an intact D-V axis and a VNC, but there was a loss of eyes, brain lobes, and foregut tissue. At later cleavage stages, BMP exposure resulted in the formation of a third ectopic eye, brain lobe, and foregut lobe. Only with exposure during late cleavage did a clear trunk phenotype appear in which ventral midline structures were lost. The loss of the ventral midline also resulted in a collapse of the longitudinal connectives of VNC into the ventral midline; however, most of the neural tissue in the VNC was still present, and the overall D-V axis did not appear to be greatly affected.

## Material and Methods

### Animal care and embryo collection

Adults of *Capitella teleta* Blake et al. (2009) were cultured in glass finger bowls with 32–34 ppt artificial sea water (ASW) at 19 °C and fed with sieved mud (Grassle and Grassle, 1976; Meyer et al., 2015; Seaver et al., 2005). Embryos of the correct stage were collected from females with broods produced from mating dishes by combining males and females after keeping them separate for 3–5 days. Embryos were collected 12–16 hours (h) after assembling the mating dish or by exposing the dish to light for 6+ h, then combining them 5 h prior to collecting zygotes (Lanza and Seaver, 2020b). Embryos and larvae, except where otherwise noted, were raised in ASW with 50 μg/mL penicillin and 60 μg/mL streptomycin (ASW+PS) at room temperature (RT, ~21 °C). ASW+PS was changed once or twice daily. Prior to fixation, all larvae were relaxed in 1:1 ASW:0.37 M MgCl_2_ for 5–10 min.

Embryos were staged by the birth of all four micromeres for a specific quartet except for the 4^th^ quartet, where only birth of 4d was monitored due to the difficulty in seeing 4a–4c being born. Thus 1^st^-quartet (1q), 2^nd^-quartet (2q), and 3^rd^-quartet (3q) embryos had micromeres 1a–1d, 2a– 2d, and 3a–3d, respectively. 4^th^-quartet (4q) animals were incubated in BMP or mock solution (see below) starting ~40 min after the appearance of micromere 4d.

### Incubation in BMP protein

Ct-BMP2/4 protein (Kenny et al., 2014) has a 42% sequence identity and 58% sequence similarity with the recombinant zebrafish BMP4 protein (R&D Systems, cat. 1128-BM-010) used; the sequence alignment was performed by T-Coffee (Di Tommaso et al., 2011). Embryos of the desired stage were isolated from a single brood and washed several times in ASW+PS. Healthy embryos—without any signs of tissue damage—were separated into 0.2% gelatin-coated, four-well culture dishes (Nunc, BioExpress cat. 144444) in 400 μL ASW+PS. For exogenous BMP treatment, animals were incubated in 250 ng/mL recombinant zebrafish BMP4 protein (R&D Systems, cat. 1128-BM-010) in 0.08% bovine serum albumin (BSA) and 0.05 mM HCl in ASW+PS for varying lengths of time (Fig. 1). Pilot experiments showed that 250 ng/mL and 500 ng/mL produced similar phenotypes, so 250 ng/mL was used for all subsequent experiments. BMP4 protein was stored as 5 or 10 μL aliquots at a stock concentration of 20 μg/mL in 4 mM HCl + 0.1% BSA at −80 °C. The two control groups included: 1) 400 μL ASW+PS (ASW), and 2) 5 μL 0.1% BSA in 4 mM HCl (mock) in 400 *μL* of ASW+PS. Animals were raised at RT until stage 6 (Seaver et al., 2005), and solutions were changed every 12 h for continuous (cont) BMP experiments. Initial pilot data showed that changing BMP every 24 h produced a less severe phenotype than 6 h or 12 h protein changes (e.g. for midline loss and VNC collapse), so 12 h changes were used. Prior to fixation, cont-treated animals were washed several times with ASW+PS to remove residual BMP or mock solutions. For pulse experiments, ASW+PS with BMP protein was added at a specific stage (Fig. 1); at the end of the pulse (1 h or 12 h), animals were washed 3–4 times in ASW+PS to remove the BMP protein (or mock solution for controls) and animals were raised to stage 6 in ASW+PS.

**Figure 1.**
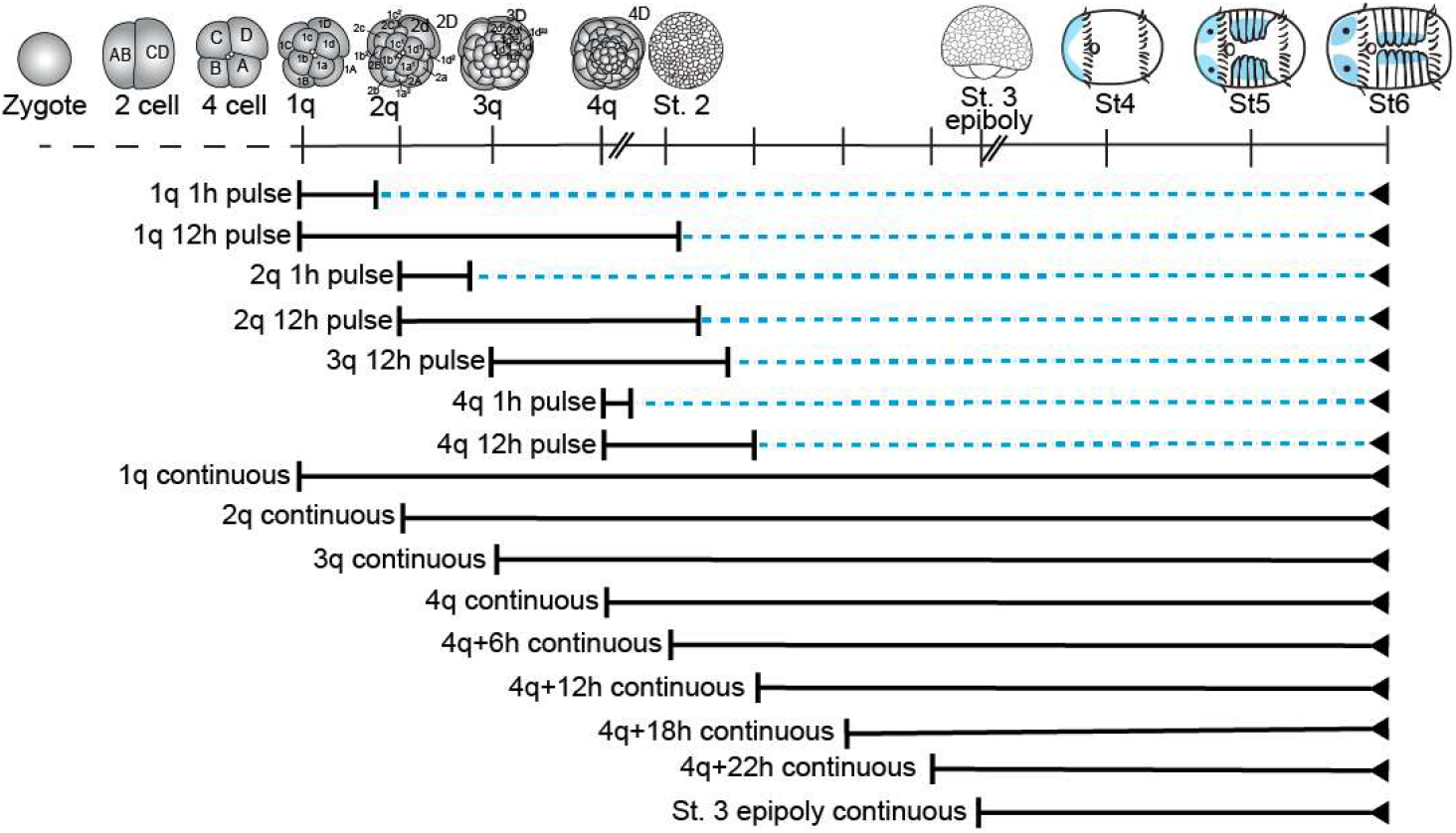
List of exogenous BMP4 treatments performed. Stage timings are approximate. Blue dashed line: Animals raised in pen/strep ASW. Blue tissue in stages 4-6: neural tissue.

### Isolation of *C. teleta* BMP pathway gene homologs

Total RNA was extracted from mixed stage 1–9 embryos and larvae using the RNA Trizol extraction protocol (Molecular Research Center, Inc.) or the RNeasy Mini Kit (Qiagen cat. 74104) paired with the QIAshredder columns (Qiagen cat. 79656). Reverse transcription reactions were conducted using the SMARTer RACE kit (Clontech cat. 634859) or High capacity cDNA Reverse Transcription kit (Applied Biosciences cat. 4368814). *Capitella teleta* has one *chordin-like* (*Ct-chrdl*) and one *bmp5-8* homolog (Kenny et al., 2014). A 940 bp fragment of *Ct-chrdl* (JGI PID224618) coding sequence was amplified by PCR using gene-specific primers: *Ct-chrdl:* 5’-ACACGAATGGAGCAGTAACTTG and 5’-AGCGCTCTGTCAGTAATTTTCA and nested primers 5’-AGAAACGCACAAGAGCCAAC and 5’-AGAAACGCACAAGAGCCAAC. A 1445 bp fragment of *Ct-bmp5-8* (JGI PID172350) was amplified with primers: 5’-ATGCTCGCTGCTTTCGCCG and 5’-CAGCTTCGTTGGCAGCAAT. PCR products were TA-cloned into the pGEM-T Easy vector (Promega) or PCRII vector (Invitrogen) and sequenced. These gene fragments were used as a template to generate DIG-labeled, anti-sense RNA probes using MegaScript SP6 or T7 transcription kit (ThermoFisher Scientific) for whole mount *in situ* hybridization (WMISH). The probe *Ct-elav1* has been previously described (Meyer and Seaver, 2009).

### Whole Mount *In situ* Hybridization

WMISH was conducted as described previously (Seaver et al. 2001). Briefly, all WMISH fixations were done in 4% paraformaldehyde (PFA, stock 32% PFA ampules from Electron Microscopy Sciences, cat. 15714) in ASW for 6 h–overnight at 4°C. After fixation, animals were serially dehydrated in methanol and stored at −20 °C. Animals were hybridized for a minimum of 72 hours at 65 °C with 1 ng/ μl of each probe. Spatiotemporal RNA localization was observed using an NBT/BCIP color reaction. The color reaction was stopped using 3 washes of PBS + 0.1% Tween-20. After WMISH, animals were labeled with 0.1 μg/ml Hoechst 33342 (Sigma-Aldrich, cat. B2261), cleared in 80% glycerol in PBS, and mounted on slides for DIC and fluorescent imaging.

### Staining and antibody labeling

Immunolabeling was carried out as in Meyer et al. (2015). Briefly, animals were fixed for 30 min with 4% PFA in ASW at RT, rinsed with PBT (PBS + 0.1% Triton-X 100), blocked in 5 or 10% heat-inactivated goat serum in PBT (block) and incubated in primary antibody in block overnight at 4 °C. Secondary antibodies in block were incubated overnight at 4 °C, then animals were thoroughly washed with PBT and cleared and mounted in 80% glycerol in PBS. All washes and exchanges were done in RainX-coated glass spot dishes. Primary antibodies used were as follows: 1:800 rabbit anti-serotonin (5HT; Sigma-Aldrich, cat. S5545), 1:10 mouse anti-Pax (clone DP311, (Davis et al., 2005), 1:800 rabbit anti-FMRFamide (Immunostar, cat. 20091), 1:20 mouse anti-Futsch (clone 22C10, Developmental Studies Hybridoma Bank), 1:800 mouse anti-acetylated tubulin (clone 6-11B-1, Sigma, cat. T6793), 1:50 mouse anti-Synapsin (clone 3C11, Developmental Studies Hybridoma Bank), and 1:800 rabbit anti-phosphorylated-SMAD1/5/8 (clone 41D10, Cell Signaling Technologies). Secondary antibodies used were as follows: 1:2000 goat anti-mouse F(ab’)2 Alexa488 (Sigma-Aldrich, cat. F8521) and 1:1000 sheep anti-rabbit F(ab’)2 Cy3 (Sigma-Aldrich, cat. C2306). F-actin and DNA staining were performed by incubating the embryos in 1:100 BODIPY FL-Phallacidin (Life Technologies, cat. B607; stock concentration 200 Units/mL in methanol), 0.1 μg/mL Hoechst 33342 (Sigma-Aldrich, cat. B2261), and/or 1:1000 ToPRO3-iodide in PBS according to Meyer et al. (2015, 2010).

#### pSMAD immunolabeling

4-cell embryos were separated into six treatments: 1h ASW, 1h BMP, 2h ASW, 2h BMP, 1h BMP + 1h ASW chase, and 1h ASW +1h BMP. 250 ng/mL recombinant BMP4 was used as previously described. The egg membrane was first softened with 3 min of 1:1 solution of 1 M sucrose and 0.25 M sodium then the embryos were fixed for 15–20 min in 4% PFA at RT, and all other steps were carried out as above. Animals were labeled with 1:800 anti-phosphorylated-SMAD1/5/8 (clone 41D10, Cell Signaling Technologies), BODIPY FL-Phallacidin, and Hoechst 33342 as above to detect increased levels of BMP signaling.

### DiI labeling

Pressure injection of the lipophilic DiI (1,1’-dioctadecyl-3,3,3’3’-tetramethylindocarbocyanine perchlorate, Invitrogen) has been described previously (Meyer et al., 2010; Sur et al., 2020). DiI diluted in soybean oil was injected into the 1a, 1b, 1c, or 1d micromeres using a M-152 micromanipulator (Narishige) and a PLI-90A picoinjector (Harvard Bioscience). Injections were done on a SteREO Discovery V8 Stereo microscope with an X-Cite fluorescent light source and a KL 1500 LED light source (Zeiss). After microinjection, normal cell division of all micromeres relative to uninjected controls was monitored for a few hours. 250 ng/mL BMP4 protein in ASW+PS was added after birth of the 4d micromere for 12 h at RT; embryos were then washed 3–4 times in ASW+PS and raised until stage 6 as described above. Animals were fixed, stained with Hoechst and Phallacidin in PBS, and cleared and imaged in 80% glycerol in PBS as previously described (Meyer et al. 2010).

### Western blot

Embryos were collected in 1.5 mL microcentrifuge tubes and briefly centrifuged in a minifuge to concentrate animals at the bottom of the tube. As much ASW as possible was removed, and animals were flash frozen in liquid nitrogen and stored at −80°C until use. Tissue was lysed until no visible pieces remained just prior to use in 15 μL of lysis buffer (50 mM Tris pH 8.0, 150 mM NaCl, 1% Triton X-100) with a pestle (Electron Microscopy Sciences, 64788) and a LabGen 7B homogenizer (ColePalmer, EW-04727-09). 13 μL of SDS solution (100 mM Tris-HCl pH 6.8, 4% w/v SDS, 20% v/v glycerol, 0.05% v/v β-mercaptoethanol, and 0.025% bromophenol blue) was added and samples were denatured at 100 °C for 5 min. Samples (n = 80 embryos / lane) were run on a precast SDS-PAGE gel (4-20% MP TGX Stain-Free Gel, Bio-Rad 4568093), with a Precision Plus Protein WesternC Stds ladder (Bio-Rad 1610376). Total protein was visualized with a stain-free viewer (Bio-Rad EZ imager) to confirm an even amount of protein in each lane. Protein was transferred to a polyvinylidene difluoride (PVDF) membrane (Bio-Rad, 1704274) with a Trans-Blot Turbo Transfer System (Bio-Rad) for 30 mins at RT. Blots were blocked for 1 h at RT with 5% bovine serum albumin in TBST (20 mM Tris pH 7.5, 150 mM NaCl, 0.1% Tween-20), incubated overnight at 4°C with a 1:1000 dilution of rabbit anti-phosphorylated-SMAD1/5/8 (clone 41D10, Cell Signaling Technologies) in block, washed with TBST, and then incubated overnight in a 1:2000 dilution of horseradish peroxidase-linked anti-rabbit IgG secondary antibody (A24537 Life Technologies) in block. The protein was visualized with Clarity ECL (Bio-Rad, 1705060) on a LAS-4000 (GE) imager.

### Microscopy and figure preparation

Images were taken using DIC optics on an AxioImager M2 microscope (Zeiss) with an 18.0-megapixel EOS Rebel T2i digital camera (Canon) for WMISH animals or an AxioCam MRm rev.3 camera (Zeiss) with Zen Blue software (Zeiss) for antibody labeled animals. DiI-labeled animals were imaged using an Zeiss Apotome.2 to produce optical sections. Animals for confocal laser scanning microscopy were cleared and mounted in SlowFade Gold (Life Technologies, cat. S36936) and imaged using a TCS SP5-X (Leica). DIC images taken at different focal planes were merged with Helicon focus 7 (Helicon); Different channels and z-stacks of fluorescent images were merged using Zen Blue (Zeiss). Images were edited for contrast, brightness and color balance (WMISH only) using Adobe Photoshop CC (Adobe). Figure panels were assembled with Adobe Illustrator CC (Adobe).

### Phenotypic scoring

#### Eye pigment

Larvae were scored for the presence/absence and location of the orange larval eye pigment cell shortly following fixation as the eye pigment fades within a week after fixation. They were scored in PBS using a Zeiss Stereo Lumar V.12 stereomicroscope. Three-eyed animals were scored in two categories: “stacked”, where the third larval eye pigment cell was in close proximity to either the left or right larval eye pigment cells, or “centered”, where the third larval eye pigment cell was closer to the ventral midline than to either the left or right larval eye pigment cells.

#### Brain scoring

Anterior z-stacked images were scored for the locations of 5HT^+^ neuronal cell bodies relative to the position of the prototroch and foregut. In Fiji (Schindelin et al., 2012), the episphere was divided into 8 sections (octets: dorsal, left, ventral, right, and the 4 intervening regions), and each 5HT^+^ soma was scored for position within the octets. The total number and rough position (i.e. left brain, right brain or ventral/ectopic brain) of acetylated tubulin positive sensory cells (sc^ac+^ cells) (acetylated tubulin positive sensory cells, Amiel et al., 2013) was also scored using anterior, z-stacked images.

#### Measurements

All measurements were made on st 6 larvae using Fiji/ImageJ2 (Rueden et al., 2017; Schindelin et al., 2012) Animal length was measured from prototroch to telotroch to control for episphere size variation.

Foregut measurements were made using Hoechst nuclear stain from a ventral view, and measurements were taken at the same focal plane as the ventral mesodermal bands in order to maximize foregut area. Total foregut area was measured including the interior space. Foregut shape was also scored as normal, small, or three-lobed.

The total width of *Ct-chrdl* expression was measured from dorsal images at segment 4. In animals expressing *Ct-bmp5-8*, the width of the dorsal midline tissue between the two bands of *Ct-bmp5-8* expression was measured from a dorsal view but was highly variable between segments, even within a single animal. The total width of *Ct-elav1* expression in the VNC was measured from left to right from ventral images at segment 4. To calculate the amount of neural tissue, the width of the ventral midline tissue not expressing *Ct-elav1* was also measured, and this width was subtracted from the total *Ct-elav1* width.

### Statistics and graphing

All statistics were performed in R/RStudio 1.2.5 (R Core team, 2014; RStudio Team, 2012), and all graphs were created using the ggplot2 package (H. Wickham, 2009) and polished with Adobe Illustrator CC (Adobe). Model testing was used in each case to determine the appropriate covariables to analyze in each ANOVA; the R package rcompanion (Mangiafico, S.S., 2015) was used, and the model with the lowest AIC was chosen.

Mean circular orientation of 5HT^+^ neurons in the episphere was analyzed using a model-based maximum likelihood approach to test multiple possible orientations (Fitak and Johnsen, 2017). The best model to fit the data was chosen using a likelihood ratio test and default settings in the CircMLE package.

## Results

### Recombinant BMP4 protein in *C. teleta* increases pSMAD1/5/8 levels

Incubating embryos and larvae in recombinant BMP protein is an effective way to test BMP function during different developmental time windows. To confirm the ability of recombinant zebrafish BMP4 protein to activate the BMP pathway in *C. teleta*, we assayed levels of phosphorylated Ct-SMAD1/5/8 (pSMAD1/5/8) in BMP-treated animals by immunostaining and western blot analysis using a cross-reactive antibody against the C-terminus of mammalian pSMAD1/5 (41D10, Cell Signaling Technologies). The transcription factor SMAD1/5/8 is phosphorylated by BMP receptor 1 (BMPR1) once bound by a BMP ligand. pSMAD1/5/8 then binds SMAD4 and is transported into the nucleus where it affects gene transcription (Gámez et al., 2013). High pSMAD levels are an indicator of active BMP signaling.

Adding recombinant BMP4 to 4-cell embryos for 1 h (n = 15) or 2 h (n = 13) resulted in visible pSMAD1/5/8 in the nuclei of all non-dividing cells relative to ASW controls (Fig. 2A-D’), and the signal was not noticeably attenuated after washing BMP protein out and incubating for another 1 h in ASW (n = 18; Fig. 2E, E’). We did not detect pSMAD1/5/8 in control embryos in ASW (1 h, n = 15; 2 h, n = 9). pSMAD1/5/8 was generally not visible in the nucleus of cells that were dividing (M phase), making it difficult to confirm that pSMAD1/5/8 was present in all cells after incubation in recombinant BMP.

**Figure 2.**
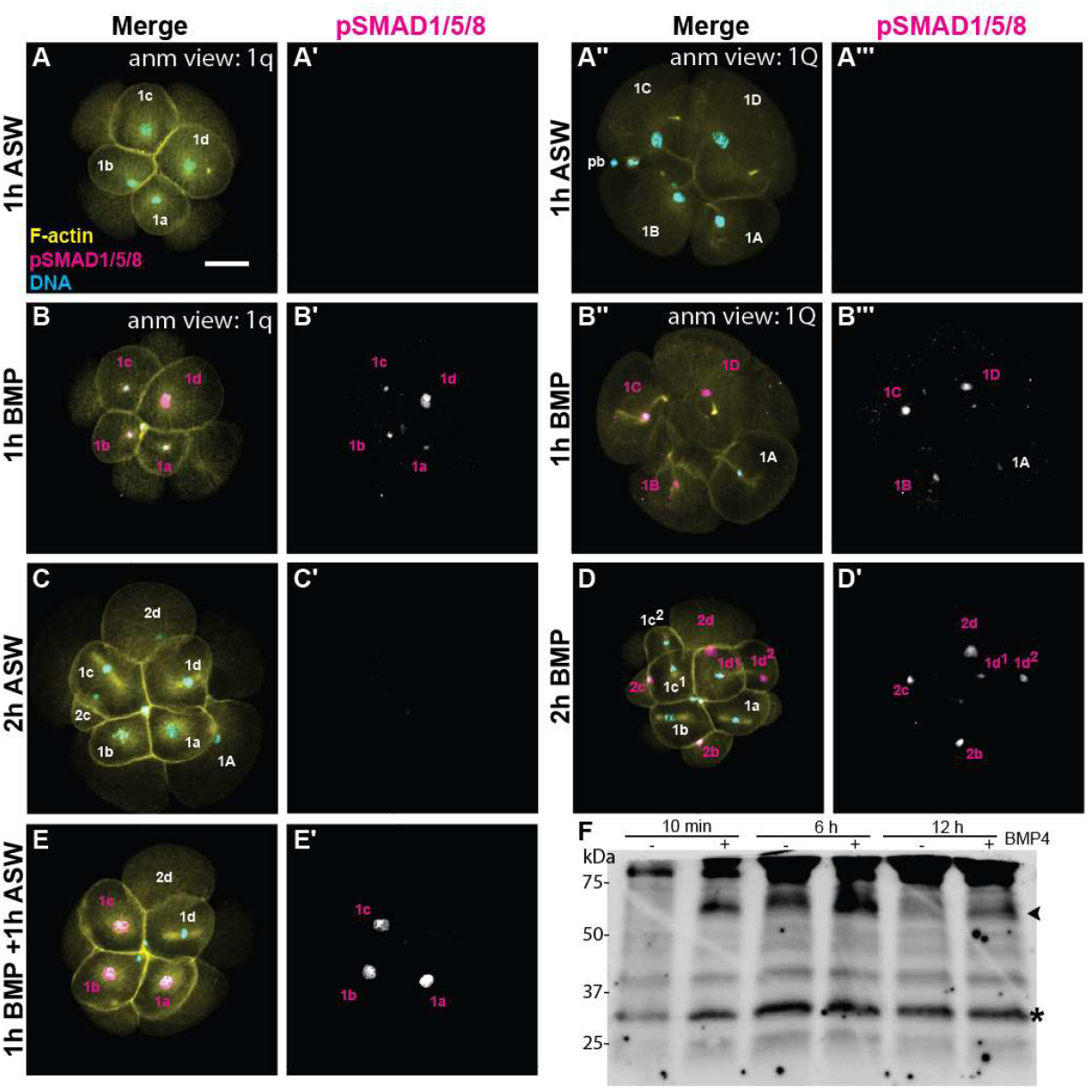
Increase in pSMAD1/5/8 after BMP exposure. (A-E) z-stacked images of immunostaining, A) 1h ASW treatment (8 cell embryo). B) 1h BMP treatment (8 cell embryo). C) 2h ASW treatment (10 cell embryo). D) 2h BMP treatment (13 cell). E) 1h BMP followed by 1h ASW treatment (9 cell embryo). (A-D) merged micromeres; (A’-D’) pSMAD1/5/8 from (AD); (A”-D”) merged macromeres, (A’”-D’”) pSMAD1/5/8 from (A”-D”). yellow: F-actin; magenta: pSMAD1/5/8; cyan: DNA. F) Western blot of pSMAD1/5/8 on 2q animals with either mock (-) or BMP (+) for 10min, 6 h, or 12 h (n = 80 per lane). *, presumed pSMAD1/5/8; arrowhead, unknown band, see text.

BMP treatment of cleavage-stage embryos with the 2^nd^-quartet of micromeres (2q; n = 80/treatment) for 10 min, 6 h, and 12 h resulted in an increase in two separate bands at ~33 and 60 kDa on the western blot relative to mock treatment of 2q embryos (Fig. 2F). The same pattern of pSMAD bands and the same pattern of up-regulation was observed on repeated western blots after treating *C. teleta* cleavage-stage embryos with BMP4 protein for various amounts of time (data not shown).

*Capitella teleta* has one SMAD1/5/8 homolog (Kenny et al., 2014), and based on genomic sequence and expressed mRNA, *Ct-SMAD1/5/8* is missing the conserved start codon (Lanza and Seaver, 2020a) found in other *SMAD 1/5/8* homologs including in the annelid *Helobdella robusta* and the molluscs *Crassostrea gigas* and *Crepidula fornicata* (Fig. 3). We cloned and sequenced *Ct-SMAD1/5/8* from *C. teleta* mixed stage embryonic and larval mRNA, and confirmed the lack of a conserved start codon. *Ct-smad1/5/8* is approximately 920 bp (vs 1395 bp in *H. robusta*) and lacks the MH1 domain, which is involved in DNA binding (Heldin et al., 1997), as well as the nuclear localization signal and one of two nuclear export signals (Xiao et al., 2003). This should produce a protein that is 381 aa or ~33 kDa. Full-length SMAD1/5/8 homologs in other animals, which do not contain this truncation, are predicted to be ~51 kDa (Fig. 3). We interpret the lower 33 kDa band on our westerns as phosphorylated Ct-SMAD1/5/8 using the start codon found in the unconserved region between MH1 and MH2. The upper band (~60 kDa) may represent an alternative upstream start site, possibly a non-canonical start site (Kearse and Wilusz, 2017), which would explain the highly conserved 5’ SMAD1/5/8 sequence (Fig. 3). Another possibility is that this upper band represents a second SMAD1/5/8 homolog in *C. teleta* that has yet to be identified and the conservation in 5’ sequence for the truncated *Ct-SMAD1/5/8* is due to a very recent loss of its start codon. Taken together, we interpret the increase in the bands on the pSMAD1/5/8 western and the immunohistological data as evidence that recombinant BMP4 protein activates the BMP signaling pathway in *C. teleta*, resulting in phosphorylation of a Ct-SMAD1/5/8 homolog that translocates to the nucleus.

**Figure 3.**
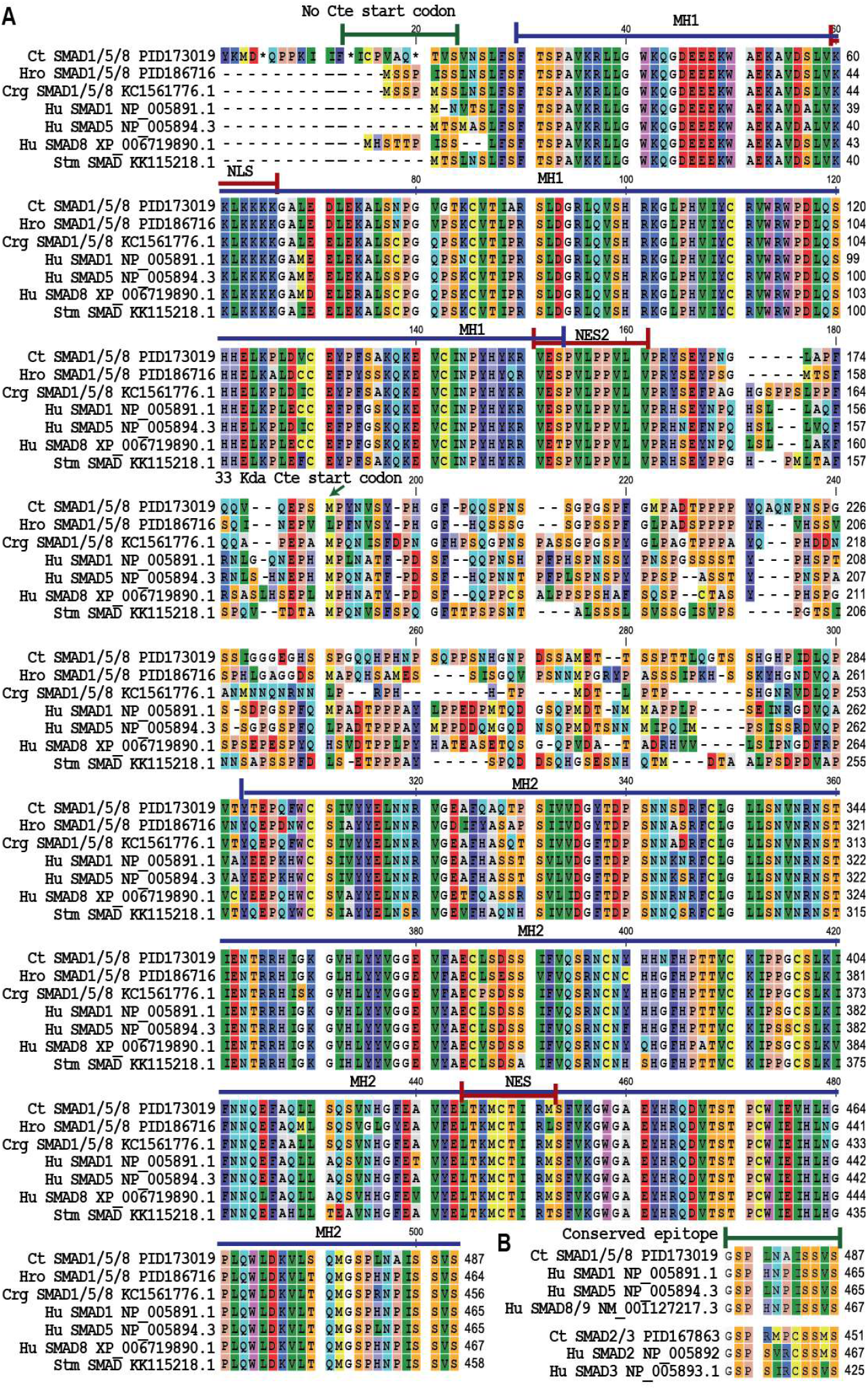
SMAD Alignments of *C. teleta* A) SMAD1/5/8 showing the loss of the conserved start codon (~18aa), the next start codon is at 225 aa. B) SMAD2/3 and SMAD1/5/8 alignments showing the conserved C-terminus where the pSMAD1/5/8 antibody 41D10 and the pSMAD2/3 antibody D27F4 bind. Species are as follows: *Ct-Capitella teleta*, Hro – *Helobdella robusta*, Crg – *Crassostrea gigas*, Hu – human, and the spider Stm - *Stegodyphus mimosarum*. Protein domains: MH1, MAD homology 1; MH2, MAD homology 2; NES, nuclear export signal; NES2, nuclear export signal 2;NLS, nuclear localization signal.

### BMP treatment in *C. teleta* does not affect overall A-P and D-V axis formation

To assess the effect of ectopic BMP signaling at different times during embryogenesis, *C. teleta* embryos were incubated in exogenous BMP4 protein for 12-hours (pulse) or continuously (cont) starting as early as birth of the 1^st^-quartet of micromeres (referred to as “1q”), i.e. 8-cell embryos, through mid-larval stages, usually stage 6 (Fig. 1). Additional starting times included: once the 2^nd^-quartet micromeres were born (2q), once the 3^rd^-quartet micromeres were born (3q), once 4d was born (4q), and at various timepoints after birth of 4d until the beginning of gastrulation (Fig. 1). Gross morphology was assayed by DIC microscopy, and, in general, features along both the dorsal-ventral (D-V) and anterior-posterior (A-P) axes were correctly localized relative to one another in stage 6 (Fig. 4) and stage 7 larvae. For the D-V axis, features examined included the foregut, chaetae, VNC, and neurotroch (ventral midline ciliary band), and features examined along the A-P axis included the brain, prototroch, foregut, and telotroch. Overall, our data suggest that neither the D-V nor A-P axes were greatly affected by BMP treatments at any of the time windows examined.

**Figure 4.**
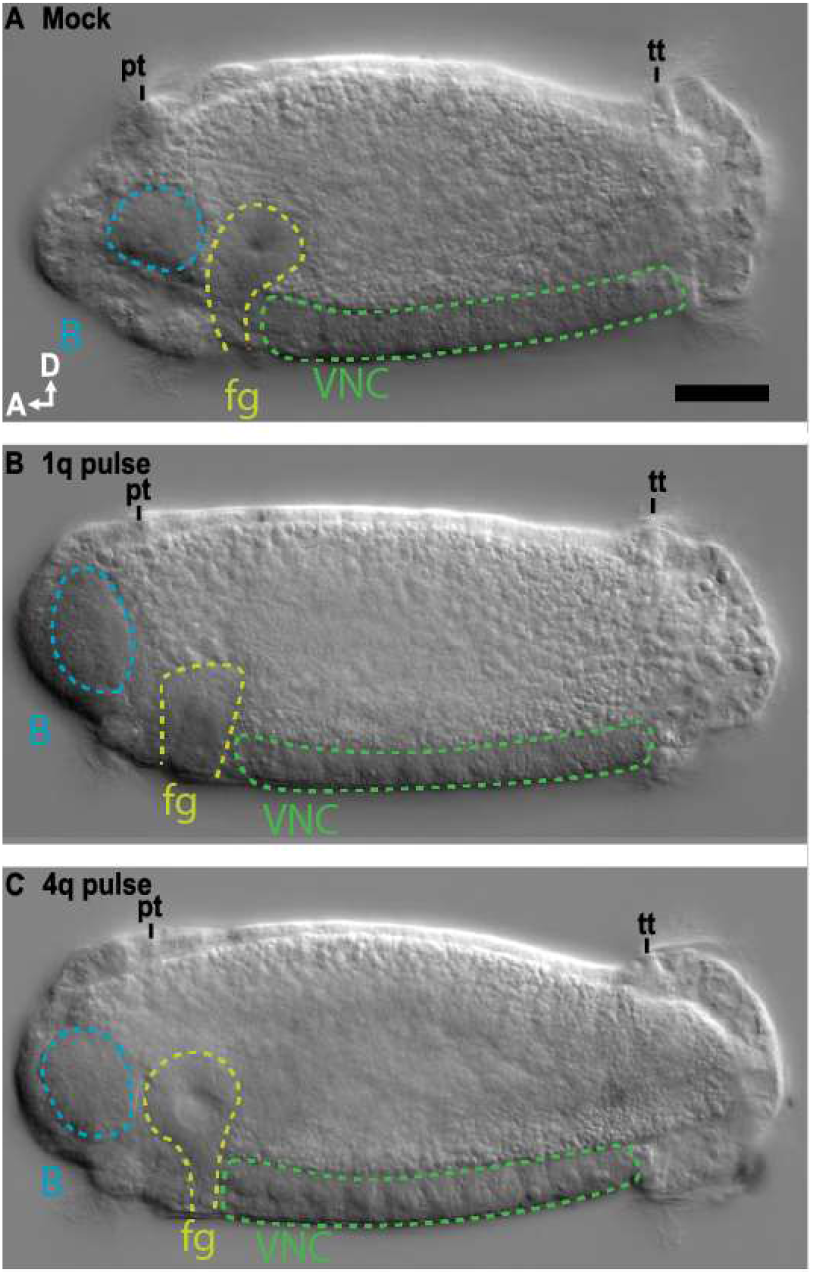
Left lateral z-stacked DIC images showing D-V axis formation in A) Mock, B) 1q pulse, and C) 4q pulse animals. brain, B (blue); foregut, fg (yellow); prototroch, pt; telotroch, telotroch; VNC (green), A, anterior; D, dorsal, scale bar = 50 μm.

To further assess overall development following BMP treatment, the number of trunk segments were counted using a nuclear stain. The number of trunk segments in BMP-treated animals was significantly reduced compared to controls, even when controlling for animal length (2-way ANOVA treatment|animal length, F_treatment_ = 33, df = 3, p <0.001). There were significantly fewer segments produced with earlier BMP treatment (1q vs 4q), but not with a longer time of BMP treatment (1q pulse vs. 1q cont) (Tukey HSD, p < 0.01). Mock animals had a mean of 12 ± 1 SD segments, 1q pulse and 1q cont animals had 9 ± 1 SD segments each, and 4q pulse had 11 ± 2 SD segments. Adding BMP4 protein reduced the number of segments produced in a timingdependent manner, which could be indicative of a slight developmental delay. This is further supported by a slight delay observed for the onset of swimming behavior in the BMP-treated animals (not scored).

### Expression of the dorsal, ectodermal markers *chordin-like* and *bmp5-8* is spatially shifted after BMP treatment

Although the formation of the D-V axis did not appear to be greatly affected in *C. teleta* larvae treated with BMP protein, we examined the expression of markers in the dorsal ectoderm since BMP signaling has been reported to pattern D-V fates within the ectoderm in two other annelids (Denes et al., 2007; Kuo et al., 2012). Chordin-like homologs in other animals antagonize BMP signaling (Branam et al., 2010), and *Ct-chordin-like* (*Ct-chrdl*) is dorsally expressed in the ectoderm starting at stage 5 (Fig. 5A) as is *Ct-bmp5-8* (Webster et al., in prep). *Ct-chrdl* expression was significantly wider (laterally) in BMP-treated animals when compared to mock animals, even when animal width and length were considered (ANOVA, F_treatment_ = 16.7, df = 3, p < 0.001, Fig. 5B, compare with 5A). All three treatments examined, 1q cont, 3q cont, and 3q pulse animals, had a wider domain of dorsal *Ct-chrdl* expression than mock animals (Tukey HSD, p < 0.07). There was no observed difference in the anterior-posterior extent of *Ct-chrdl* (Fig. 5A, B).

**Figure 5.**
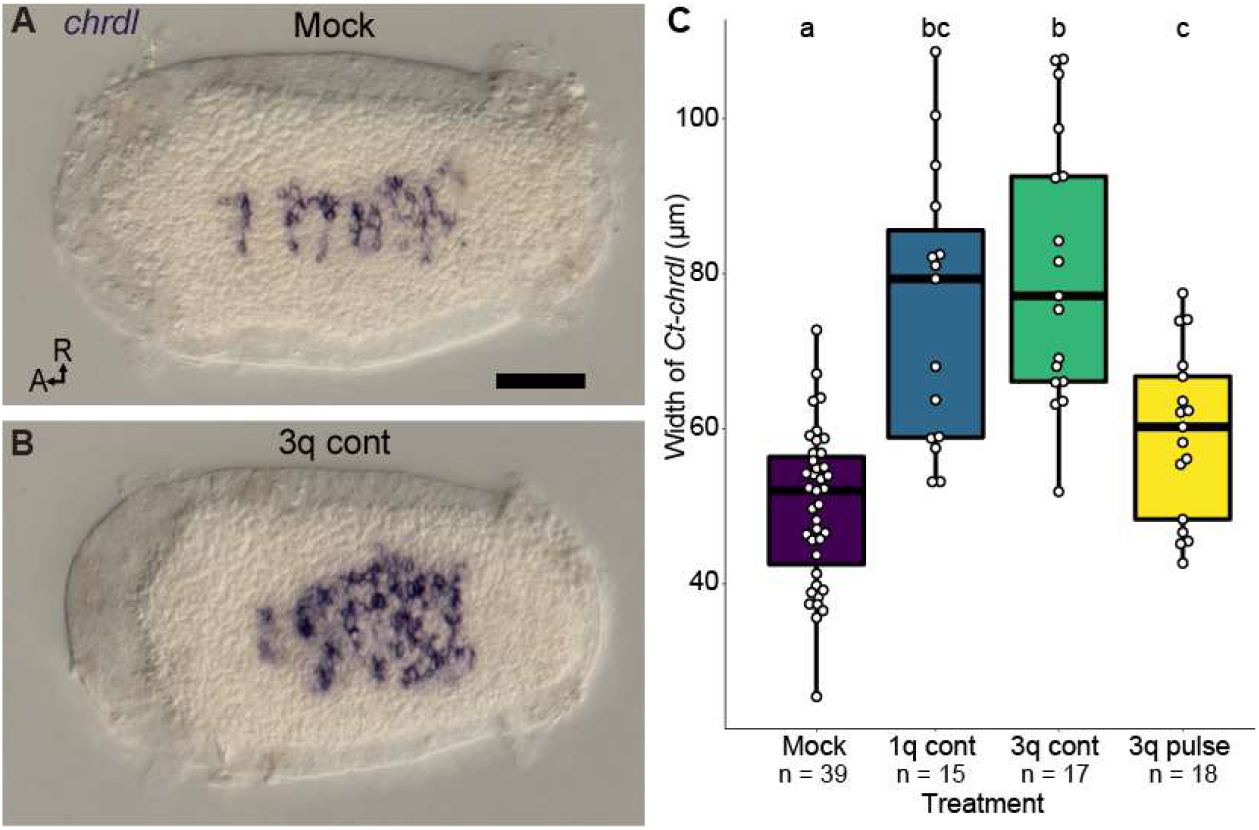
*Ct-chrdl* expression. (A-B) Dorsal z-stack of *Ct-chrdl* mRNA expression in (A) Mock and (B) 3q cont BMP treated animals. C) Boxplot showing the significant effect of BMP treatment on the width of Ct-chrdl expression, letters indicate significance groups. A, anterior; R, right side; scale bar = 50 μm.

*Ct-bmp5-8* is expressed as two stripes in the dorsal-lateral ectoderm starting at stage 5, on either side of the dorsal *Ct-chrdl* stripe (Webster et al., in prep). In agreement with the increased domain of *Ct-chdl* expression, the dorsal boundaries of the *Ct-bmp5-8* stripes were shifted more ventrally (i.e. the width between left and right bands of expression was greater) with a 1q or 3q pulse of BMP (data not shown).

### BMP treatment during early cleavage stages does not block brain or VNC formation

To look at the effect of BMP treatment on neural tissue, we first examined formation of the brain and VNC using the marker *Ct-elav1* (*CapI-elav1*) (Fig. 6), which is expressed throughout the nervous system in post-mitotic neurons (Meyer and Seaver, 2009; Sur et al., 2020, 2017). In the episphere after BMP treatment, *Ct-elav1* was expressed in the brain, but the tissue organization was grossly affected with varying location and number of brain lobes (Fig. 6B, C, D). Furthermore, the episphere of BMP-treated animals appeared smaller (e.g. Fig. 4), with the *Ct-elav1^+^* tissue comprising a larger proportion (data not shown). 1q pulse and 1q cont animals generally showed a disorganized pattern of *Ct-elav1* (Fig. 6B, D), while 4q pulse animals generally had three brain lobes based on nuclear staining, all of which expressed *Ct-elav1* (Fig. 6C; further discussed below).

**Figure 6.**
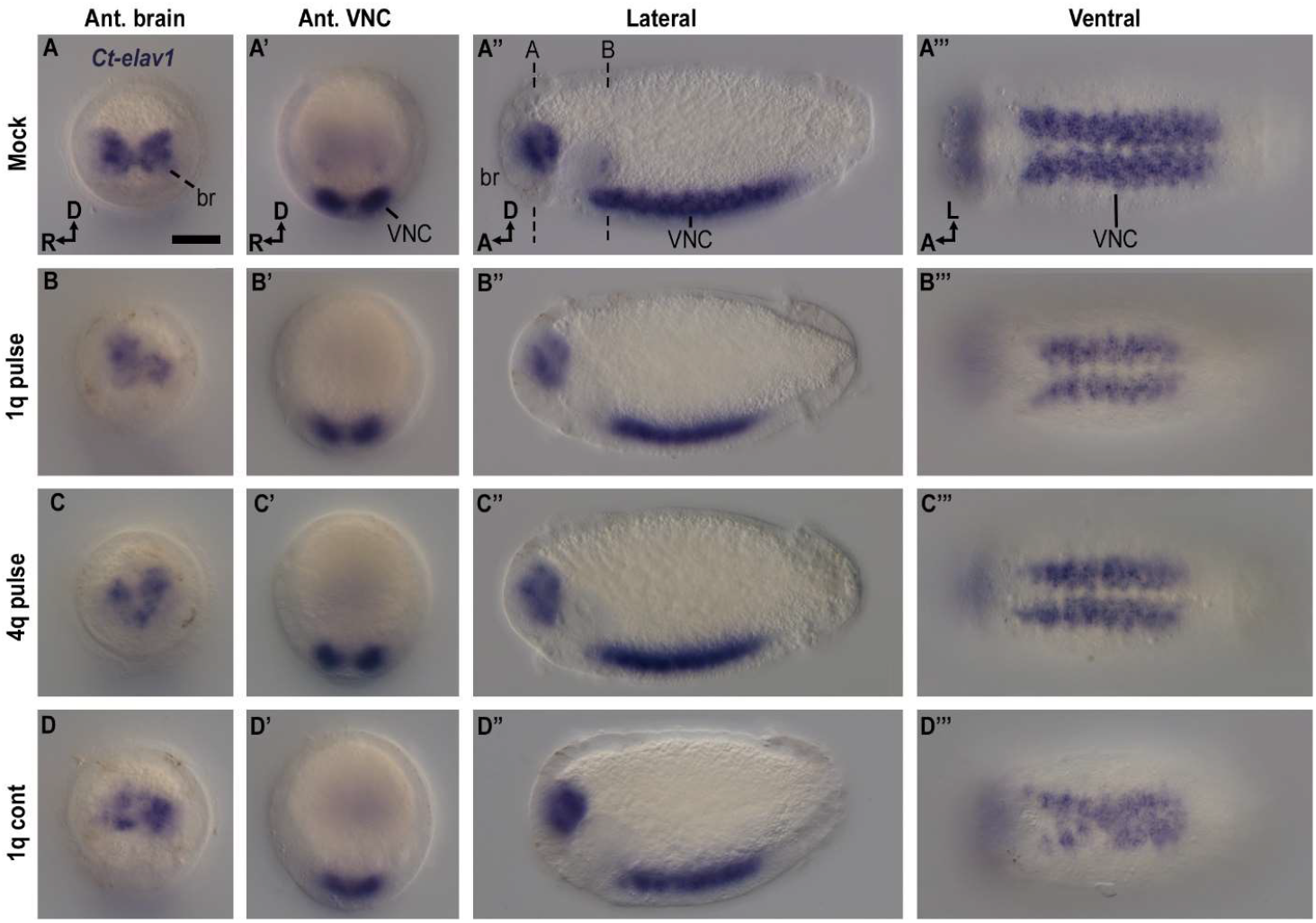
*Ct-Elav1* expression in BMP-treated animals. (A-D) Mock (n = 37), dashed lines in (C) represent focal depths of (A) and (B). (E-H) 1q pulse (n = 36). (I-L) 4q pulse (n = 36). (M-P) 1q cont (n = 45) showing collapse of some hemiganglia expressing Elav1. Scale bar = 50 μm; br: brain, VNC: ventral nerve cord, A, anterior; D, dorsal, L. left;

Overall, *Ct-elav1* expression in the VNC of animals treated with a pulse of BMP starting at 1q or 4q was similar to that in wild-type and mock animals, with bilaterally-symmetric expression in the ganglia on either side of the ventral midline (Fig. 6A’”-D’”). However, in animals treated continuously with BMP from 1q to larval stage 6, *Ct-elav1* was expressed in the ganglia of the VNC, but the ventral midline gap was missing between the left and right hemiganglia in each segment (Fig. 6D’”). While BMP treatment did not appear to block formation of the VNC, and the neural tissue was still ventrally-positioned in the trunk (e.g. Fig. 6A’-D’), we also quantified whether BMP treatment affected the amount of neural tissue within a segment. This was done by measuring segment 4—where the most severe collapse was evident in continuously-treated animals. The total width from left to right of the *Ct-elav1* expression domain was measured, and any *Ct-elav1*^-^ midline gap was excluded to approximate the total amount of neural tissue in segment 4. A pulse of BMP starting at 1q or 4q did not significantly affect the total width of *Ct-elav1* expression minus the midline gap relative to mock-treated animals; mock (n = 37), 67.0 ± 4.6 μm (SD); 1q pulse (n = 36), 60.0 ± 6.7 μm (SD); 4q pulse (n = 36), 61.7 ± 6.1 μm (SD). In contrast, cont 1q BMP treatment had a significant effect on the width of tissue expressing *Ct-elav1* (ANOVA, F_treatment_ = 78, df = 3, p < 0.001; Tukey’s HSD, p < 0.001, Fig. 6), where 1q cont animals had a significantly narrower expression domain in segment 4 (Mean width: 1q cont (n = 45), 50.3 ± 13.6 μm (SD); Fig. 6P). This pattern also held true for the width of *Ct-elav1* expression, *including* the midline gap (data not shown), and was not affected by animal size (model testing; prototroch–telotroch length). Taken together, these data suggest that pulses of BMP at 1q and 4q do not change the amount of neural tissue in the VNC, at least in segment 4, while continuous BMP treatment at 1q does reduce the amount of neural tissue in segment 4 (~25% reduction), even when the loss of the non-neural midline tissue is taken into account.

We also found that BMP treatment had a significant effect on the A-P length of tissue expressing *Ct-elav1* even when the prototroch-telotroch length was accounted for (2-way ANOVA treatment| prototroch-telotroch length, F_treatment_ = 58.7, df = 3, p < 0.001). All treatments produced a significantly shorter *Ct-elav1* expression region (i.e. VNC) than in mock animals (Figure 6), which probably relates to the reduced total number of segments seen with BMP treatment.

### BMP treatment during late cleavage stages resulted in a loss of ventral midline structures and collapse of the VNC

To further explore the architecture of the VNC after BMP treatment, we examined nuclei, neurites (anti-acetylated-Tubulin labeling), and serotonergic neurons (anti-5HT labeling) in late stage 6/early stage 7 larvae. BMP did not have a strong effect on formation of the VNC in 1q pulse, 4q pulse, or stage 3 cont treatments (Fig. 7). The ganglia, longitudinal connectives, commissures, and 5HT^+^ neurons in the VNC and the peripheral nerves in each segment were clearly visible (Fig. 7 and data not shown). There appeared to be a slight developmental delay in the number of 5HT^+^ longitudinal connectives in the 1q and 4q pulse animals (compare 7B, C with Fig. 7A), and the longitudinal connectives appeared disorganized in larvae treated from stage 3 to late stage 6 (Fig. 7D).

**Figure 7.**
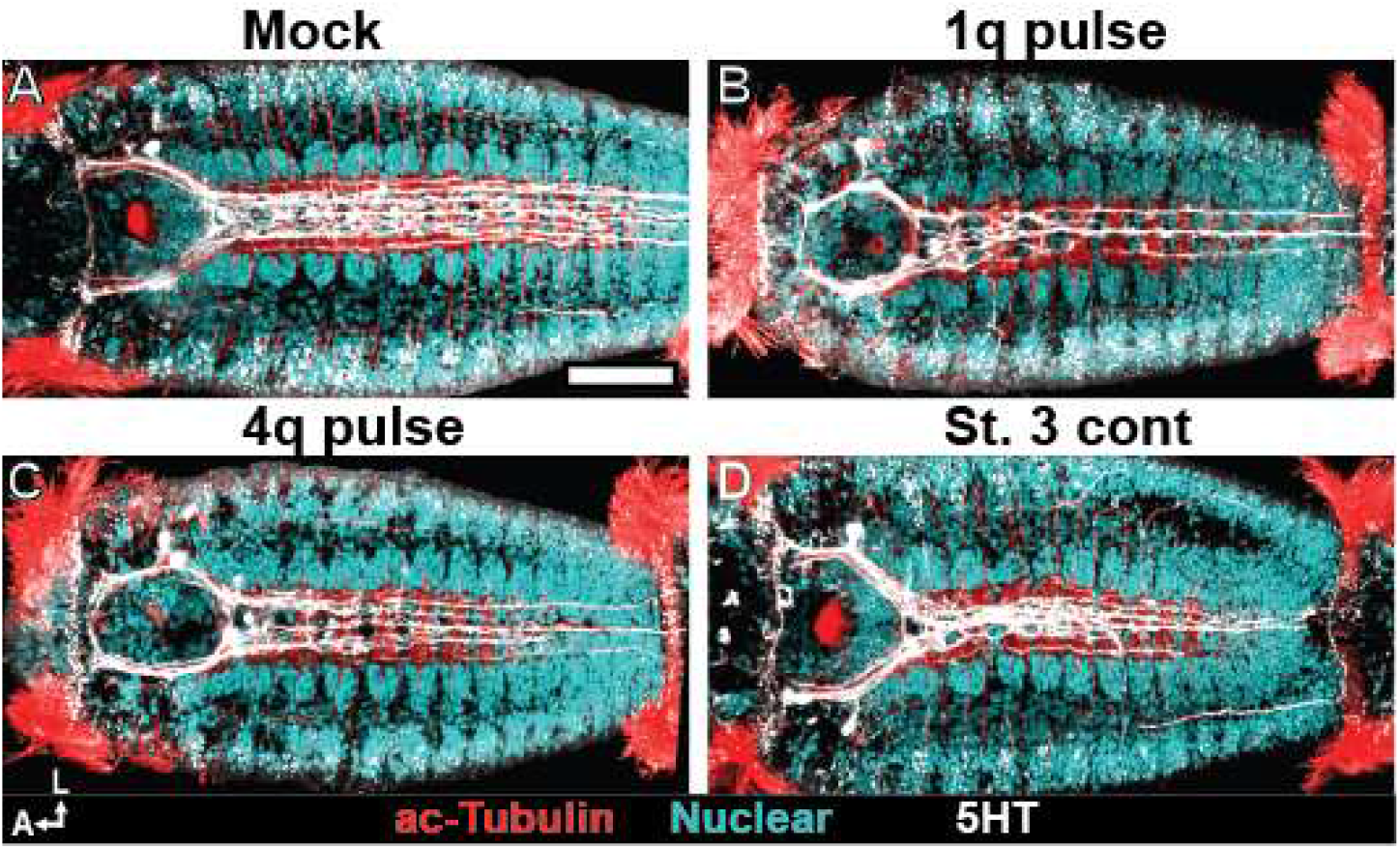
Ventral z-stack of the serotonergic (5ht, white), and anti-acetylated-Tubulin staining (red) in the VNC with a nuclear stain (blue). Only minor disorganization was seen in 1q pulse (B), 4q pulse (C), or st 3 cont (D) animals compared to mock (A). A, anterior; L, left side; scale bar = 50 μm.

Since continuous BMP treatment starting at 1q resulted in expression of *Ct-elav1* across the ventral midline at stage 6, we further examined formation of the neurotroch and the architecture of the VNC using Hoechst (nuclei) and cross-reactive antibodies against acetylated-Tubulin (cilia and neurites), serotonin, FMRF, Synapsin, and Pax (Figs. 8 and data not shown). Animals were treated continuously with BMP protein starting at 1q, 2q, 3q, 4q, 4q+6h, 4q+12h, 4q+18h, 4q+22h (just prior to the onset of gastrulation), and stage 3 (onset of gastrulation) until stage 6. The severity of midline phenotypes varied from complete loss of the midline along the anterior-posterior extent of the trunk to mild loss of the midline in a few segments. In the most severely affected animals, the cilia of the neurotroch were not detectable (Fig. 8B), while in more mild cases, patches of neurotroch cilia were present, often towards the posterior-end of the trunk. In all animals treated with BMP, the cilia around the mouth were present despite loss of the neurotroch (e.g. Fig. 8B, C). It is worth noting that the ciliary cells around the mouth arise from a different blastomere lineage than the rest of the ciliary cells in the neurotroch (3c and 3d versus 2d^2^ and 2d^12^, respectively) (Meyer et al., 2010; Meyer and Seaver, 2010). Loss of the neurotroch was accompanied by a collapse towards the midline of the connectives of the VNC. In the most severe cases of VNC collapse, all five main connectives (acetylated-Tubulin^+^ and Synapsin^+^), including the three 5HT^+^ connectives and the two FMRF^+^ connectives, formed a single track along the midline (Fig. 8B’, E’ and data not shown). Furthermore, the left and right hemiganglia appeared continuous across the midline in all segments (Fig. 8E), and subsets of Pax^+^ neural cells that are normally bilaterally-symmetric on either side of the midline in each segment (Fig. 8D”, arrowheads) were present in the ventral midline (Fig. 8E”, arrowheads). We also saw a range of intermediate phenotypes, including BMP-treated animals with no midline abnormality (data not shown) and animals with a ‘mild’ phenotype consisting of partial VNC collapse, hemiganglia intercalation, and patchy reduction of neurotroch cilia in a few segments (Fig. 8C-C’, F-F”). We did not observe VNC collapse or loss of the neurotroch in mock-treated animals or in animals treated with a 12 h pulse of BMP protein starting at 1q, 2q, 3q or 4q.

**Figure 8.**
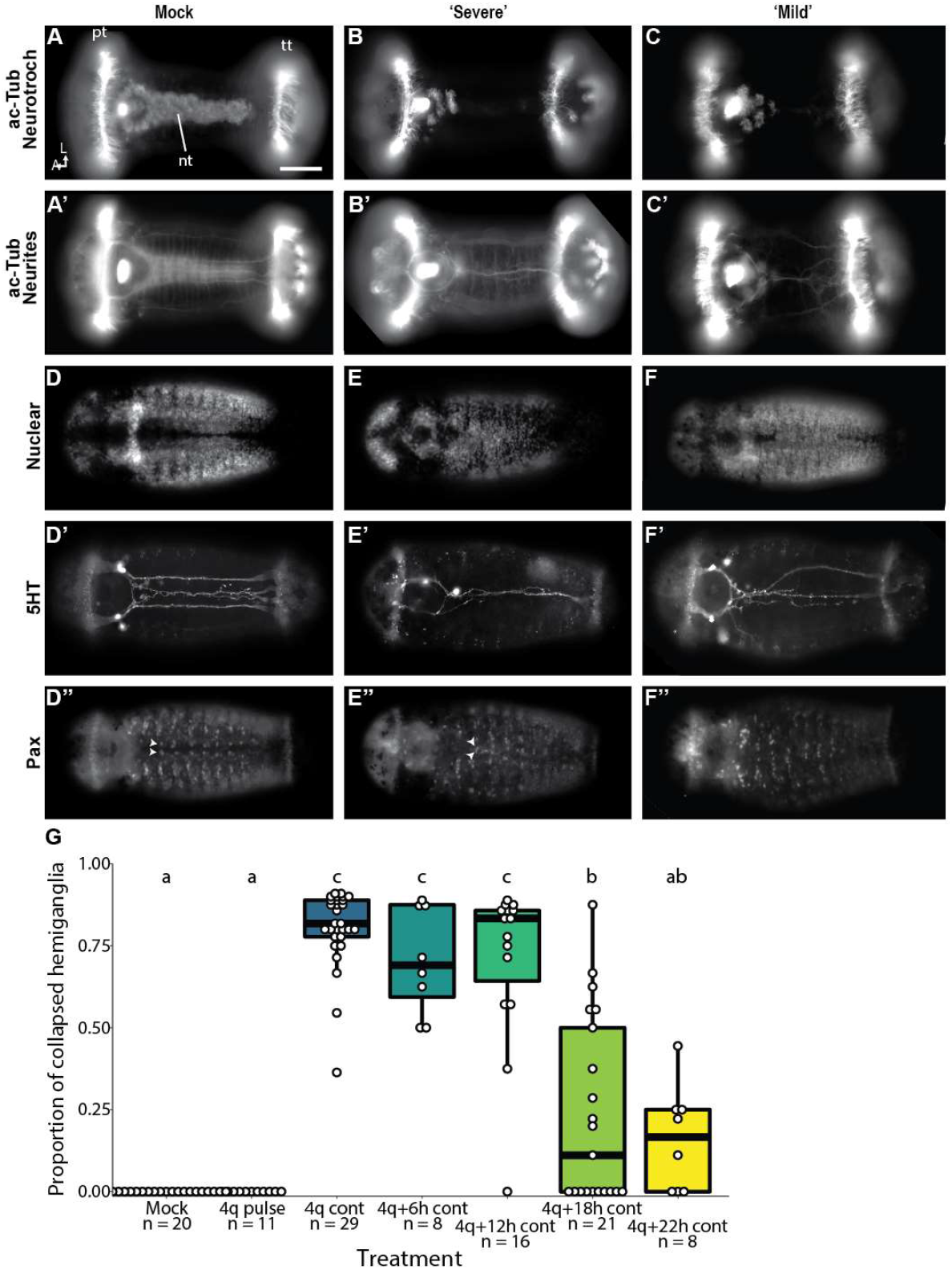
Trunk neural phenotype in cont BMP-treated (Z-stack). (A,D) A mock animals showing wild-type phenotype. (B,E) 4q+22h continuous animals showing a severe VNC collapse phenotype. (C,F) 4q+22h continuous animals showing a mild VNC collapse. (A-C) acetylated-Tubulin showing neurotroch cilia, (A’-C’) acetylated-Tubulin in the VNC. (D-F) Hoechst nuclear stain. (D’-F’) Serotonin (D’’-F’’) Pax, arrowhead: normally bilaterally-symmetric cells on either side of the midline in each segment. Figure panels with the same letter represent the same animal. A, anterior, L, left side, nt, neurotroch; pt, prototroch; tt, telotroch; VNC, ventral nerve cord; scale bar = 50 μm (G) Boxplot showing the significant effect of treatment on proportion of segments with collapsed hemiganglia, letters indicate significance groups.

To determine the timing critical to cause VNC collapse, we quantified the proportion of segments with collapsed hemiganglia from cont BMP treatment starting at different times. The degree of VNC collapse varied significantly with treatment (Fig. 8P, Two-way ANOVA treatment | total segment number, F_treatment_ = 70, df = 6, p < 0.001). VNC collapse was most penetrant when BMP protein was added just after the birth of the 4d micromere (4q) or 6 or 12 h afterwards (4q+6h and 4q+12h, respectively) and changed every 12 h until stage 6 (cont treatment). The frequency of VNC collapse gradually decreased in animals continuously incubated in BMP protein starting at 4q+18h and 4q+22h, which is just prior to the onset of gastrulation. Adding BMP protein continuously starting at early cleavage stages (1q, 2q, 3q, or 4q) also showed a highly penetrant VNC collapse phenotype (data not shown). Taken together, these data suggest that the critical time window to cause a complete VNC collapse is 12–18 h after birth of the 4d micromere, with some collapse seen with treatment up to the onset of gastrulation (4q+22h). Continuous exposure to BMP protein from gastrulation to stage 6 was not sufficient to cause a VNC collapse. A pulse of BMP protein encompassing 12–18 h after birth of the 4d micromere may be sufficient to cause a VNC collapse phenotype; however, this was not tested.

The number and location of segments with either loss of neurotroch cilia or collapsed hemiganglia were scored to look for correlation between these two phenotypes. Segments showing collapsed hemiganglia were significantly more likely to have lost neurotroch cilia (Yang’s Chi-Squared test for clustered binary paired data, X^2^= 9.19, df = 1, p = 0.002), so further analysis only examined hemiganglia collapse. The same trend was evident when examining collapse of the longitudinal nerve tracks in the VNC, but these were highly variable and difficult to quantify. The VNC collapse and coincident loss of the neurotroch strongly suggests that the VNC collapse is due to a loss of the ventral midline itself. Two homologs of ‘midline markers’, *Ct-sim1* (JGI PID165634) (Denes et al., 2007) and *Ct-admp* (JGI PID184506) (Kenny et al., 2014; Scimone et al., 2014) were cloned to confirm midline loss, but neither gene was robustly expressed in the midline of control animals at stage 6 (data not shown).

### Eye formation is affected by BMP treatment in a time-dependent manner

There are two bilaterally-symmetric larval eyes in *C. teleta*, each composed of three cells: a pigment cell, a sensory cell (SC), and a support cell (Rhode, 1993). The larval eyes degrade during metamorphosis and are replaced by two juvenile or adult eyes. The left larval eye is formed by daughters of blastomere 1a, and the right larval eye is formed by daughters of blastomere 1c (Meyer et al., 2010). BMP treatment had a striking effect on formation of the pigment cells in the larval eyes (Fisher’s exact; p < 0.001, Fig. 9). The number and position of the larval eye pigment cells in mock-treated animals (both cont and pulse treatments) were not significantly different from animals raised in ASW (p = 1.0, data not shown and Fig. 9B, E). A 12 h pulse of BMP at 1q, 2q, and 3q generally prevented formation of the larval eye pigment cells (0 eye pigment cells formed, Fig. 9A, E) giving rise to an eyeless phenotype. The 3q pulse had significantly more animals with ‘other’ eye pigment categories including: 2 larval eye pigment cells not in a wild type configuration or 1 larval eye pigment cell in various locations. A 12 h pulse of BMP at 4q resulted in 3 larval eye pigment cells; in most animals (66%), the 3^rd^ eye pigment cell was ventromedial (‘central eye’ Fig. 9C, E) while the next most common phenotype was a 3^rd^ eye pigment cell that was in close proximity to either the left or right larval eye (‘stacked eye’, 20%, Fig. 6D, E). There was no left/right bias in which side the stacked ectopic eye pigment cell was located. The proportion of wild-type eye pigment cells increased significantly and the number of 3 eyes ‘stacked’ decreased significantly in animals treated with a 12-hour BMP pulse at 4q+6h when compared to a 12-hour BMP pulse at 4q (Fisher’s exact post-hoc p < 0.001, data not shown). In animals treated with a 12-hour BMP pulse starting at 4q+12h, 4q+18h, or 4q+22h, the larval eye pigment cell phenotype was not significantly different from mock animals. The number of eye pigment cells formed in cont experiments were not significantly different from pulse experiments started at the same stage (2q and 4q cont, Fisher’s exact post-hoc p < 0.1, data not shown).

**Figure 9.**
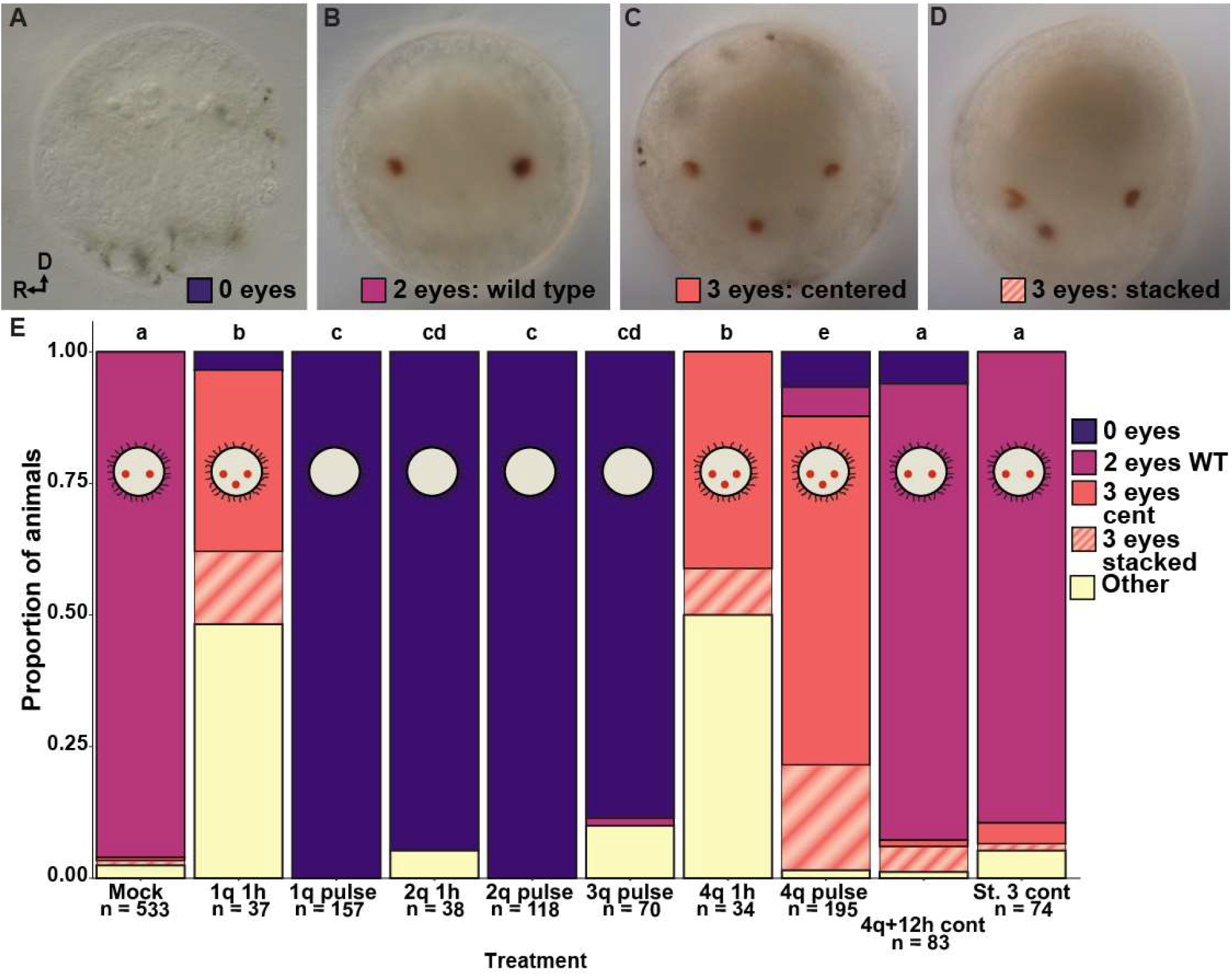
Larval eye pigment cells. A-D Most common eye (red dot) phenotypes, anterior view. A) 0 eyes (1q pulse), B) 2 wild-type eyes (1q pulse), C) 3 eyes centered (1q pulse), D) 3 eyes stacked (1q pulse). R, right side; D, dorsal. E. Proportion of animals with each eye phenotype in each BMP treatment (colors correspond to boxes in A-D, and yellow: other (see text for a description of other eye phenotypes). Letters indicate significance groups.

Shorter incubation time in BMP protein (1 h pulse instead of a 12 h pulse) was used to further explore the effect of BMP on larval eye pigment cell formation (Fig. 9E). A 1 h incubation in BMP at 1q produced a range of phenotypes, including 3 larval eye pigment cells (14/37), and a striking right-biased asymmetry where 15/17 animals with only two eye pigment cells had 2 right eyes, and 4/4 animals with one 1 eye pigment cell had only a right eye. A 1 h pulse of BMP at 2q (2q 1h) resulted in 0 larval eye pigment cells (36/38). The pattern of eye pigment cells in animals treated with BMP for 1 h at 4q (4q 1h) was very similar to the 1q 1h phenotype: many with 3 eye pigment cells (17/34) and a right-bias with 9/9 two-right eyed animals. There were also some animals with two ventral eye pigment cells (6/34).

We also examined formation of larval and juvenile eye SCs relative to the brain lobes after BMP treatment (Fig. 10), both of which are visible during larval stages using a cross-reactive antibody against the *D. melanogaster* protein Futsch (clone 22C10) (Yamaguchi and Seaver, 2013). In wild-type animals, the two larval eye SCs are located at the mediolateral edge of the left and right brain lobes (Fig. 10B, arrows). Each larval eye SC cell has a neurite that extends into the cerebral commissure as well as a bulb-shaped protrusion containing microvilli with f-actin that sits within the cup-shaped pigment cell (Fig. 10E), which is characteristic of rhabdomeric photoreceptors (Yamaguchi and Seaver, 2013). The two wild-type juvenile eye SCs are positioned next to the larval eye SCs (Fig. 10G, arrowheads), but are more superficial and have a very different morphology including a thick extension that contacts the surface of the epidermis in the episphere (Fig. 10J) and an absence of microvilli with f-actin (Yamaguchi and Seaver, 2013). In contrast to the larval eyes, juvenile SCs form from descendants of 1c and 1d (Yamaguchi et al., 2016). We quantified the number of larval and juvenile eye SCs after BMP treatment and found different patterns. Larval eye SCs followed the same pattern as the larval eye pigment cells. A 1q pulse of BMP led to 0 SCs (Fig. 10A, C), and the brain appeared to be radialized with nuclei throughout the episphere (Fig. 10C; discussed below). A 4q pulse of BMP led to 3 SCs, most of which were centered (Fig. 10A, D). Furthermore, animals with a third centered eye SC also appeared to have a third, ventromedially-positioned brain lobe (e.g. Fig. 10D; discussed below). Overall, *C. teleta* formed 0 larval eyes (pigment cells and SCs) with BMP treatments starting at 1q–3q versus 3 larval eyes with treatments starting at 4q; by 4q+12h larval eye formation was wild-type.

**Figure 10.**
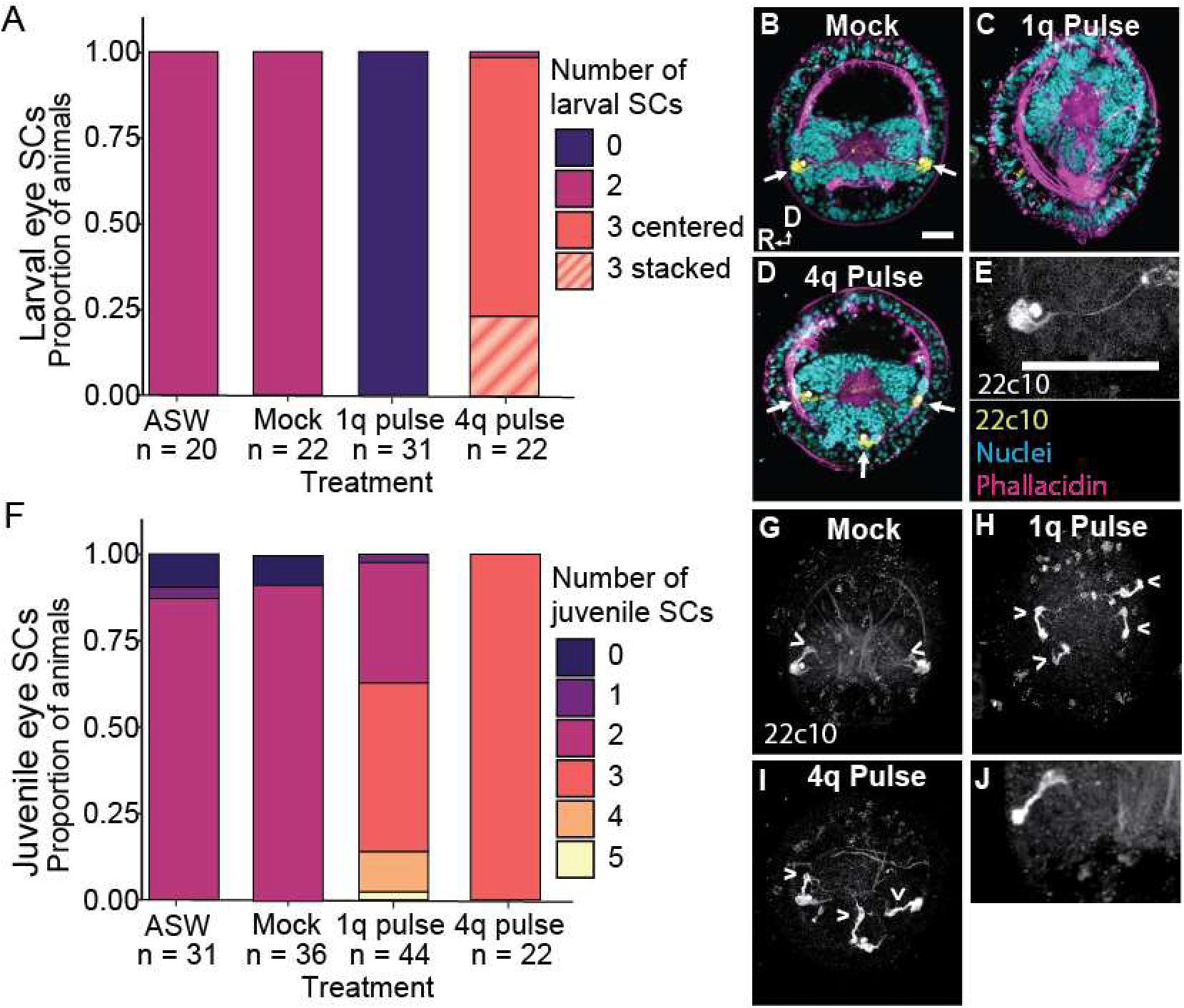
The effect of BMP treatment on larval and juveniles sensory cells (SCs). A-E) Each larval SCs was associated with an eye pigment cell. F-J) BMP increased the number of juvenile eye SCs. A) Proportion of animals with each larval SC phenotype in each BMP treatment. Letters indicate significance groups; colors are the same as in Fig. 9. B-E, G-L) Anterior z-stack of 22c10 showing (B-D) larval SCs (yellow, arrows), nuclei (teal), and muscles (magenta). E) Image of single larval SC. F) Proportion of animals with each number of juvenile SCs in each BMP treatment. G-L) juvenile SCs (arrowheads). B,G) Mock, C,H) 1q pulse, and E,I) 4q pulse animals. R, right side, D, dorsal, scale bars = 50 μm.

In contrast, the juvenile eye SCs increased in number in both 1q and 4q pulse-treated animals (Fig. 10F, H, I). In 1q pulse animals, which did not have larval eye pigment cells or SCs, there were a mean of 2.8 ± 0.8SD juvenile eye SCs and a maximum of 5 juvenile eye SCs (Fig. 10F). In 4q pulse animals, all had 3 juvenile eye SCs including an ectopic ventromedially-positioned eye SC (Fig. 10F, I). A 1q pulse of BMP increases the number of juvenile SCs, but at 4q, only 3 juvenile SCs are formed, following the pattern of larval eyes. BMP-treated *C. teleta* larvae were not raised through metamorphosis to determine if the juvenile SCs developed into functional eyes.

### Serotonergic neurons in the brain are spatially rearranged after BMP treatment

Similar to the larval and juvenile eyes in *C. teleta* larvae, serotonergic neurons in the brain arise in a bilaterally-symmetric manner, with the same number of serotonergic (5HT^+^) soma in each brain lobe at any given stage (Meyer et al., 2015). Since BMP treatment affected the number and location of the larval and juvenile eyes and also appeared to affect the arrangement and number of *Ct-elav1*^+^ brain lobes depending on the timing of treatment, we scored the position of 5HT^+^ soma in the brain to see if there were similar phenotypes (Fig. 11).

**Figure 11.**
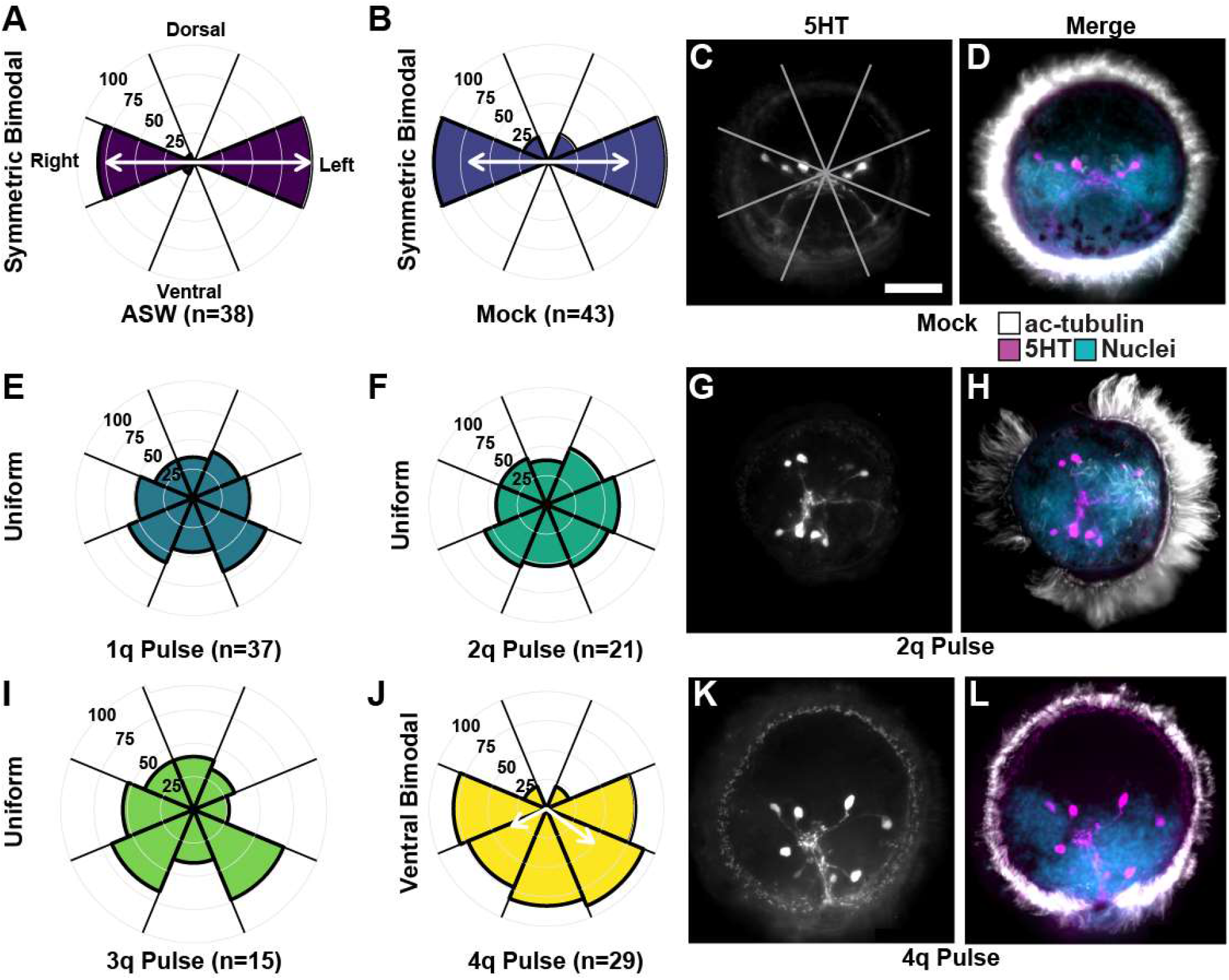
Localization of 5HT in the episphere. A-D) ASW (A) and mock (B-D) treatment animals had a symmetric bimodal distribution of 5HT cells showing a left and right brain lobe. E-I) 1q (E), 2q (F-H), and 3q (I) pulse BMP treatment animals showed a uniform distribution of 5HT cells, showing a radialized brain. J-L) 4q pulse animals showed a bimodal ventral distribution of 5HT, showing a ventralization of the brain. AB, EF, IJ) Percent of animals with 5HT cells in each region of the episphere. C,F,I) stacked immunostaining of 5HT in 3 treatments representative of the 3 brain states. D,H,I) stacked immunostaining of 5HT (magenta), ac-tubulin (white), and nuclei (cyan).White arrows: Mean vectors(s) of 5HT localization model, scale bars: 50 μm.

Most ASW and mock-treated animals (>82%) had 5HT^+^ soma that were localized only in the lateral octets of the episphere—the left and right brain lobes (Fig. 11A–D). This corresponded to a symmetric bilateral distribution (ASW: −log likelihood: 56; θ (mean angle(s)): 180° and 0°; κ (concentration): 36 and 12; mock: −log likelihood: 123; θ: 180°and 0°; κ: 8 and 8). In 1q, 2q, and 3q-treated BMP pulse animals, 5HT^+^ soma were radially localized—they were found in all regions of the episphere, which best fits a uniform distribution (1q pulse: −log likelihood: 261, 2q pulse: −log likelihood:123, 3q pulse: −log likelihood 103; Fig. 11E–I). Despite the uniform distribution, 3q pulse animals also showed a weak ventral trend. In 4q-treated, BMP pulse animals, the 5HT^+^ soma had a strong ventral distribution, which corresponded to a ventral bimodal distribution (4q pulse: - loglikelihood: 207, θ: 334° and 216°; κ: 2.2 and 4.5, Fig. 11J–L). It is important to note that there is no model to test for a trimodal distribution to statistically confirm the observed three brain lobes, a left, right, and ventral concentration of nuclei in the episphere (e.g. Figs. 6C, 10D). However, we observed 5HT^+^ soma in the third ectopic brain lobe in 4q-treated BMP pulse animals. Overall, BMP pulses at 1q–3q produce radialized 5HT^+^ cells, suggesting a radialized brain and a loss of bilateral symmetry in the episphere. At 4q, the statistically determined ventral bimodal pattern likely represents a ventral, trilobed brain.

We also quantified the total number of 5HT^+^ (serotonergic) and sc^ac+^ (acetylated tubulin positive sensory cells, Amiel et al., 2013) neurons in the brain. With a 2q pulse of BMP, we observed a complete loss of sc^ac+^ cells in the episphere (n = 30/30), whereas mock-treated animals from the same brood had a mean of 3 ± 0.6 sc^ac+^ cells on both the left and right sides for a total of ~6 sc^ac+^ cells (n = 22/22). sc^ac+^ cells were also lost after a 4q pulse (BMP n = 14/14, mock n = 15). By a 4q + 6h cont BMP treatment, most of the sc^ac+^ cells were present (4.4 ± 1.9 (SD) sc^ac+^ cells), even in animals with 3 brain lobes, but never in the ventromedial 3^rd^ brain lobe. The number of 5HT^+^ cells was significantly higher in 1q, 2q, and 3q pulse, BMP-treated animals compared to mock (ANOVA, F_treatment_ = 6.13, df = 4, p < 0.001; Tukey HSD, p < .05). The 1q, 2q, and 3q pulse, BMP-treated animals had a mean of 7.1 ± 1.0 (SD) 5HT^+^ cells (n = 44), while mock-treated animals had 5.9 ± 0.5 (SD) 5HT^+^ cells (n = 18). The number of 5HT^+^ cells was also significantly higher with a 4q BMP pulse (mean = 7.7 ± 1.0 (SD) 5HT^+^ cells, n = 15/19) than mock treated animals (mean = 6.4 ± .8 (SD) 5HT^+^ cells) (Student’s t-test, t = −3.86, df = 14, p < 0.001), but only when animals with a radial brain were removed (n = 4/19). Overall, BMP pulses at 2q and 4q causes a loss of sc^ac+^ cells, and BMP pulses starting at 1q–4q causes a small increase in 5HT^+^ cells.

### The ectopic 3^rd^ eye and brain lobe is usually derived from the 1b blastomere

To determine the origin of the third, ventromedially-positioned ectopic larval eye and brain lobe, we injected the lineage tracer DiI into blastomeres 1a, 1b, 1c, or 1d at 1q (Fig. 12). These embryos were then treated for 12 h with BMP protein starting at 4q. Our data supported previous lineage tracing where the right eye and the majority of the right brain lobe are formed by the 1c blastomere while 1b normally forms a small, ventral region of the right brain lobe and does not contribute to the two larval eyes (Meyer et al., 2010). The majority of the left brain lobe is formed by 1d, while the 1a blastomere forms the left eye and a small, ventral region of the left brain lobe. In most of the animals with DiI labeling in their larval eyes, we were able to see DiI in both the sensory cell and the surrounding pigment cell; the support cell could not be distinguished. In 3-eyed animals where blastomere 1a was labeled (n = 15), most (n = 8/15) had a centered ectopic 3^rd^ eye, n = 6/15 had a stacked ectopic 3^rd^ eye, and n = 1/15 had three ventrally-positioned eyes. All 15 animals had DiI in the left eye (Fig. 12A, magenta arrowhead) or dorsal-left eye in animals with two left-stacked eyes. We also found DiI in a second eye in n = 5/15 animals (2/8 centered, 0/2 right-stacked, 3/4 left-stacked, 0/1 ventral). All (n = 20) 1b-labeled animals with three eyes showed DiI labeling in the ectopic eye (17/20 centered, 2/20 left stacked, 1/20 right stacked). In 1b labelled animals with stacked eyes, DiI labelling was only in the ventral-most eye. Although it was difficult to confirm, some 1b-labeled animals had DiI labeling near, and possibly within their right eye; 3/10 animals with a centered phenotype and both (2/2) animals with a left-stacked phenotype. All (n = 20) 1c-labeled animals showed only DiI only in their right eye (Fig. 12C), while no (n = 17) 1d-labeled animals had DiI labeling in any eyes (Fig. 12D). Animals were imaged using a Zeiss ApoTome.2 to generate optical sections (see Methods), to confirm the presence/absence of DiI in the eyes rather than the surrounding tissue.

**Figure 12.**
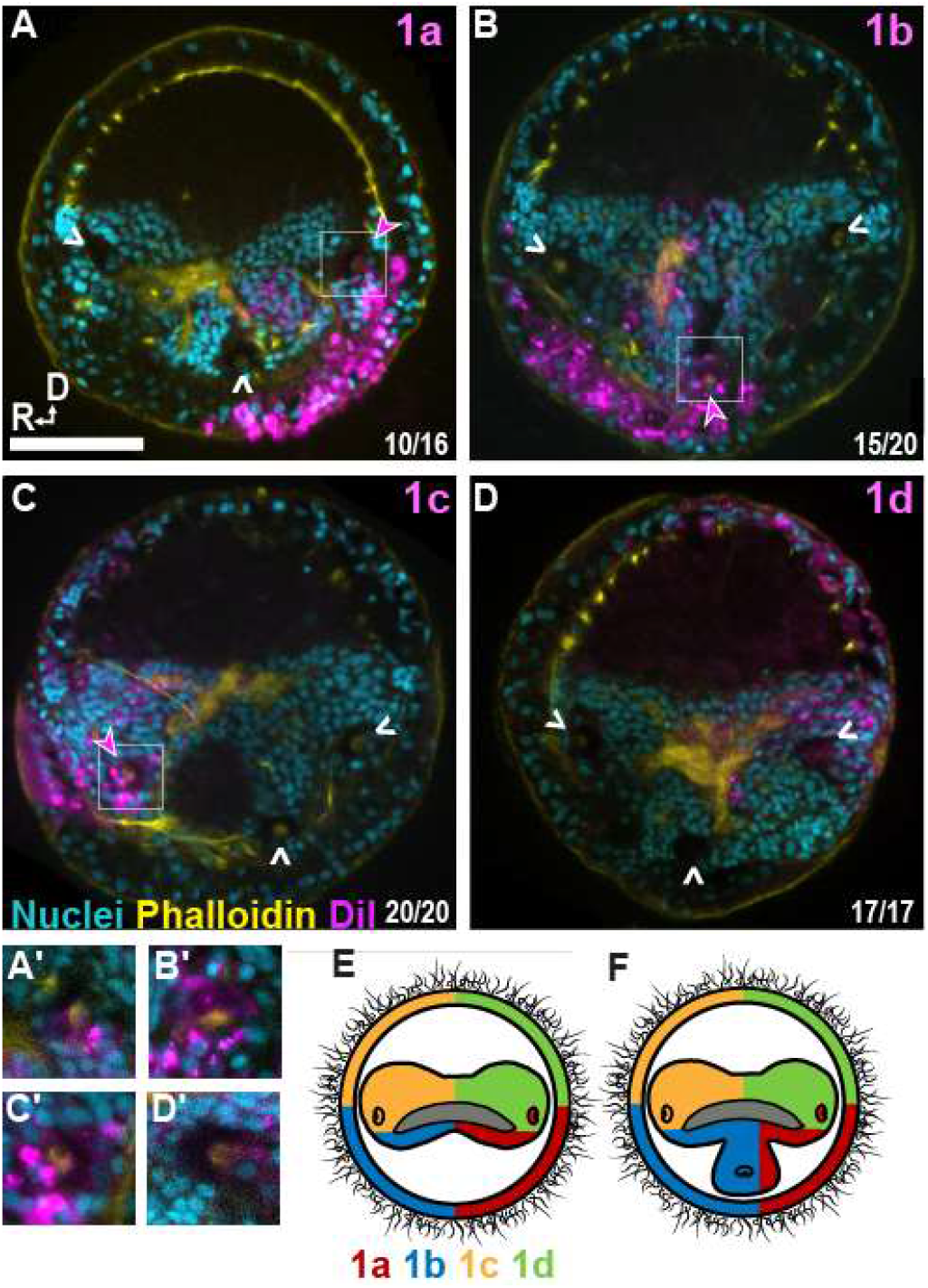
DiI localization and fate map of the eyes and brain of animals treated with BMP with a 4q pulse (Z-stack). A) 1a injected, showing contributions to the left brain lobe and eye B) 1b injected, showing contributions to the right brain lobe and the ectopic eye and brain lobe C) 1c injected, showing contributions to the right brain lobe. D) 1d injected, showing contributions to the left brain lobe. A’-C’) Inset of eye with DiI labelling (D’) Inset of eye without DiI labelling. E) Contribution of 1q micromeres to the episphere in wildtype animals. F) Predicted contribution 1q micromeres including the contribution of 1b to the third ectopic eye in BMP treated 3-eyed animals. Arrowheads, eye location; pink closed arrowhead; eye with DiI; white open arrowhead, eye without DiI; cyan, nuclei; yellow, phalloidin; magenta, DiI; scale bar = 50 μm.

It was more difficult to score DiI labeling in the brain because in some animals the ventromedial brain tissue was not morphologically distinct from either the left or right brain lobe (e.g. Fig. 12C). However, in 1b-labeled animals with a distinct, ventromedial, third brain lobe, DiI was found throughout much of the ventromedial brain lobe as well as in small, ventral portion of the right brain lobe (Fig. 12B). Animals in which 1a was labeled had DiI in a small, left portion of the ventromedial brain lobe and a small, ventral portion of the left brain lobe (Fig. 12A). Blastomeres 1c and 1d largely did not appear to contribute to the ventromedial brain lobe, as DiI was mostly found in a large, dorsal portion of the right and left respective brain lobes (see Fig. 12F for a summary).

These results suggest that in animals with a centered eye phenotype, 1b primarily forms the ectopic, ventromedial eye (Fig. 12F), although 1a may also be able to form the ventromedial eye (n = 17/17 versus n = 2/8, respectively); we did not find contributions from 1c or 1d to the ectopic, ventromedial eye. Furthermore, BMP treatment may occasionally result in a right eye that is sometimes generated by 1b instead of 1c. Finally, the ectopic, ventromedial brain lobe seems to be primarily generated by 1b with a small, left contribution from 1a (Fig. 12F).

### Foregut development is disrupted by BMP treatment in a time-dependent manner

Most of the ectodermal tissue in the episphere, including the brain and larval eyes, is generated by 1q micromeres. In order to determine if the dramatic changes after BMP treatment were confined to the episphere, we examined the foregut in the trunk, which is also ectodermal in origin, but is derived from micromeres 2a (left foregut lobe) and 2c (right foregut lobe) (Meyer et al., 2010). The foregut is normally ‘butterfly-shaped’ with two morphologically-separate left and right lobes at stage 6 (Fig. 13A, yellow dashed line) that eventually fuse at late larval stages to form the foregut. This morphology was notably affected by BMP treatment; foregut shape and ventral maximum surface area were measured to quantify that effect. All control animals (ASW and mock) showed a normal butterfly-shaped foregut (ASW: n = 12, mock: n = 37; Fig. 13A). All animals treated with BMP between 1q and 3q showed a smaller, mostly round foregut (Fig. 13B), with no left-right bias (data not shown). This included both pulse and cont treatments (1q pulse n = 50, 1q cont n = 24, 2q pulse n = 14, 3q pulse n = 28; Fig. 13B). When animals were treated at 4q, the resulting foregut had a distinct 3-lobed, triangular appearance, with what appeared to be an ectopic, posterior foregut lobe containing foregut cilia (4q pulse, n = 25/29, Fig. 13C and data not shown). The timing to produce a 3-lobed foregut was precise, and animals treated too early would generally have a small, round foregut similar to the 1q–3q phenotype (data not shown). Animals treated at 4q+6h and later had a generally normal bilobed foregut (data not shown). The foreguts of 1q, 2q, 3q, and 4q pulse, BMP-treated animals had significantly smaller areas than control animals when measured from a single focal plane coincident with the mesodermal bands (Fig. 13D, E; ANOVA [ASW, mock, 1q pulse, 2q pulse, 3q pulse], F_treatment_ = 12.0, DF = 4, p < 0.001; ANOVA [mock, 4q pulse], F_treatment_ = 5.9, DF = 1, p = 0.026). However, measurement of the total volume of the tri-lobed foreguts in 4q-treated animals could give different results as the ectopic, posterior lobe was often at different angles, making area measurement at one focal plane difficult. Foregut sizes for 1q, 2q, and 3q pulse animals were not significantly different from each other (Tukey HSD, p > 0.92). Overall, 1q, 2q, and 3q BMP pulse-treated animals had a reduced, radial foregut, while 4q pulse animals had a tri-lobed foregut (Fig. 13B, C).

**Figure 13.**
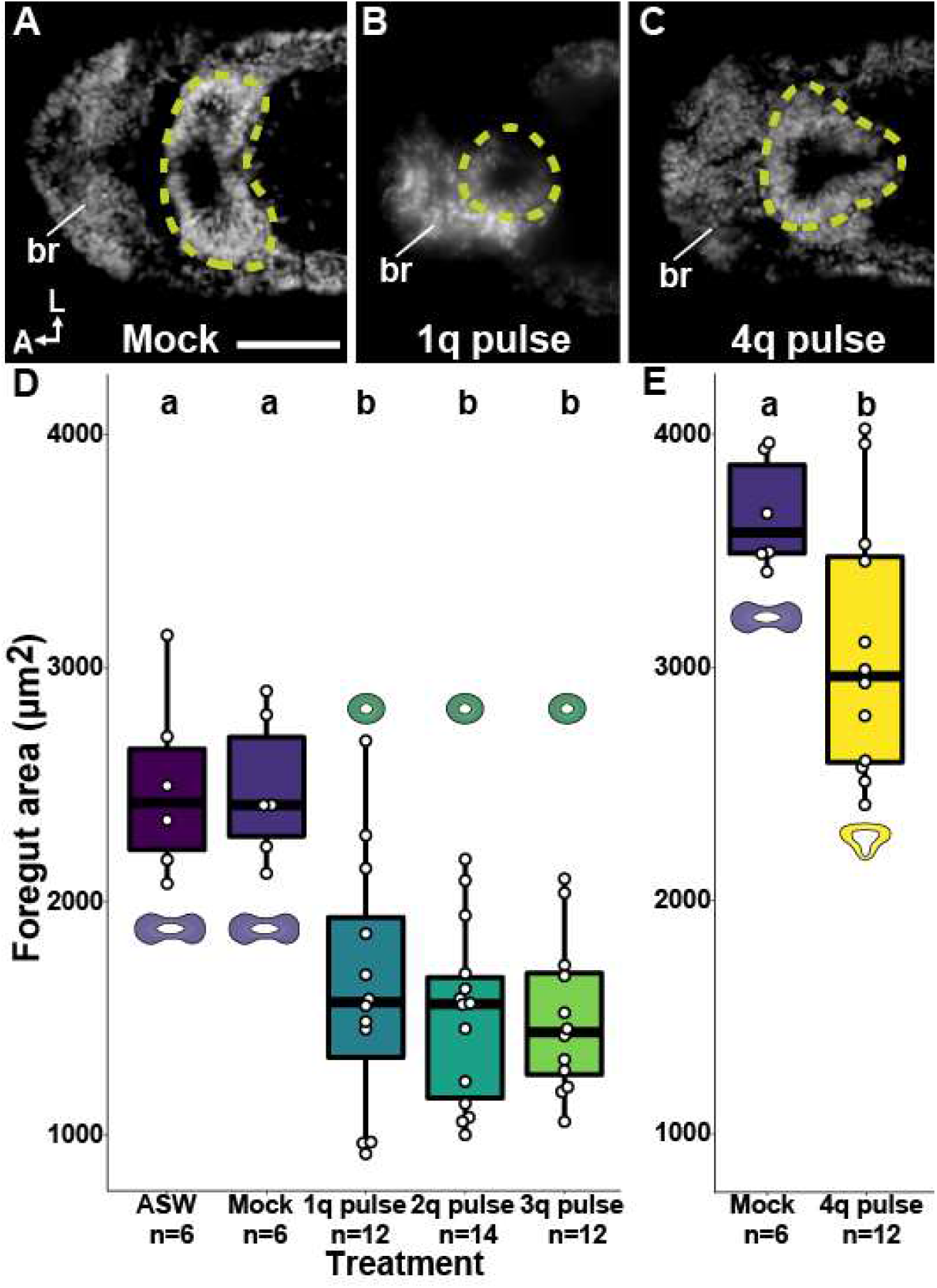
The effect of BMP treatment on foregut morphology. A) Bilobed foreguts in mock and ASW control animals. B) 1q, 2q, and 3q 12h pulse animals had small round foreguts (100%). C) Foreguts of 4q 12h pulse animals were trilobed (100%). (D-E) Total foregut area was significantly smaller in animals subjected to 1q, 2q, 3q, and 4q 12h BMP pulse than the controls. Animals in E were fixed at a later stage than animals in D and could not be compared due to significant size difference in mock animals. brain, br; foregut, yellow dashed line; scale bar: 50 μm

## Discussion

Although BMP signaling plays many roles during development, we focused on understanding its role in neural fate specification for the brain and VNC in the annelid *C. teleta*. Unlike the function of BMPs to limit the region of neuroectoderm to one side of the D-V axis in vertebrates and insects, our results demonstrate that ectopic BMP in *C. teleta* does not block formation of the brain or VNC, nor does it affect overall formation of the D-V axis. We observed a number of distinct and penetrant phenotypes that clustered into three groups depending on the time window of BMP application (Fig. 14). 1) A loss of larval eyes, a reduction of the foregut, and a radialized brain was a dominant phenotype that appears to be due to ectopic BMP treatment during the time window from around 3q to the birth of 4d. 2) A third ectopic eye, brain lobe, and foregut lobe were formed by exogenous BMP exposure between 4q and 4q+12h. 3) A loss of the ventral midline neurotroch and a collapse of the VNC was produced by ectopic BMP exposure between ~4q+12 h and 4q+22 h (i.e. during the several hours prior to the onset of gastrulation).

**Figure 14.**
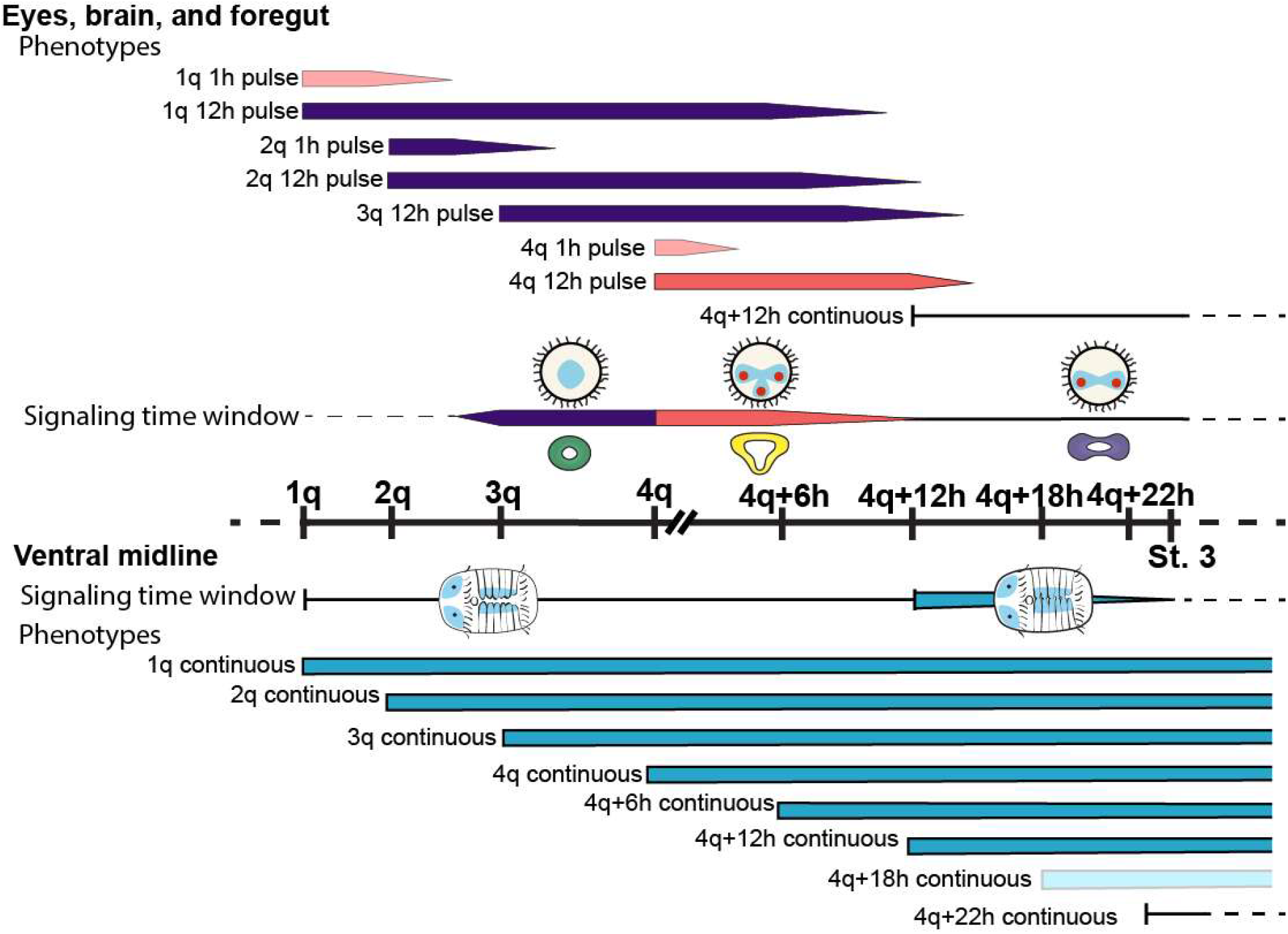
Overview of BMP treatment results and hypothesized fate switching time windows in *C. teleta*. Top: Eyes, brain, and foregut resulting phenotypes and inferred BMP signaling window. Purple: no eyes, radialized brain, small foregut. Pink: 3 eyes, 3 brain lobes, 3 foregut lobes. Bottom: Ventral midline inferred BMP signaling window and resulting phenotypes. Blue: loss of ventral midline. Black line: wild type. Lighter colors indicate less penetrant phenotype.

### BMP signaling during cleavage stages in *C. teleta* may induce quadrant-specific blastomere fates

We hypothesize that BMP signaling induces blastomere fates in a timing-dependent manner in *C. teleta*. Like many spiralians, *C. teleta* has a stereotyped cleavage pattern, where cell fates are predetermined by regulated cell division (Meyer et al., 2010). The four quadrants of the ectoderm and brain in the episphere are formed by descendants of micromeres 1a–1d; the left and right brain lobes are primarily generated by 1d and 1c with minor contributions to the ventral side of each brain lobe from 1a and 1b (Fig. 12E). The left and right larval eyes are generated by 1a and 1c, while the juvenile eye SCs form from descendants of 1c and 1d (Yamaguchi et al., 2016). The majority of the left and right lobes of the foregut, which will form the pharynx and esophagus, are generated by micromeres 2a and 2c with minor contributions to the region between the two foregut lobes from 2b, 3a, 3b, and 3d. Blastomere 2d forms the trunk ectoderm to the telotroch including the VNC and neurotroch.

The D-quadrant is of particular interest for early developmental signaling because the D-quadrant produces organizer signaling in spiralians (Henry et al., 2017). An organizer is a cell or group of cells with a large effect on development and is responsible for setting up the normal pattern of cell fates (Lambert, 2010; Nakamoto et al., 2011). In the annelid *C. teleta*, one cell at the 13- to 16-cell stage, 2d, has been identified as an axial organizer, and blastomere ablation experiments demonstrated that 2d is necessary for D-V axis formation in the episphere and for mesoderm induction in the trunk (Amiel et al., 2013). In wild-type *C. teleta* embryos, A- and C-quadrant micromeres are positioned on either side of the D-quadrant, while B-quadrant micromeres are positioned farther away from the D-quadrant, on the opposite side of the embryo. In general, the D-quadrant macromere divides first during each cleavage, followed by the C-, then B-, and finally A-quadrant macromeres.

Based on our results, we hypothesize that BMP signaling from the D-quadrant acts first on D-quadrant micromeres as they are born (with early, high, and/ or autonomous levels of BMP signaling). Subsequently, the neighboring C- and A-quadrant micromeres are born, receiving later and/or lower juxtacrine BMP signaling. The B-quadrant micromeres, which are the farthest away from the D-quadrant, may adopt their fates in the absence of BMP signaling. Although mRNA expression patterns may not represent the final localization of secreted proteins, transcripts for *Ct-bmp2/4* and the BMP antagonist *Ct-nogginA* (*Ct-nogA*) are both expressed asymmetrically along the B/D axis in early cleavage-stage embryos (Fig. 15A–C), with increased expression of both in D over B quadrants at 1q, 2q, and 3q (Lanza and Seaver, 2020b, Webster et al., in prep). The expression patterns of *Ct-bmp5-8* and the BMP antagonist *Ct-chordin-like* (*Ct-chrdl*) are more difficult to interpret as there appears to be subsets of cells in each quadrant that express both mRNAs at different stages (Fig. 15D–F).

**Figure 15.**
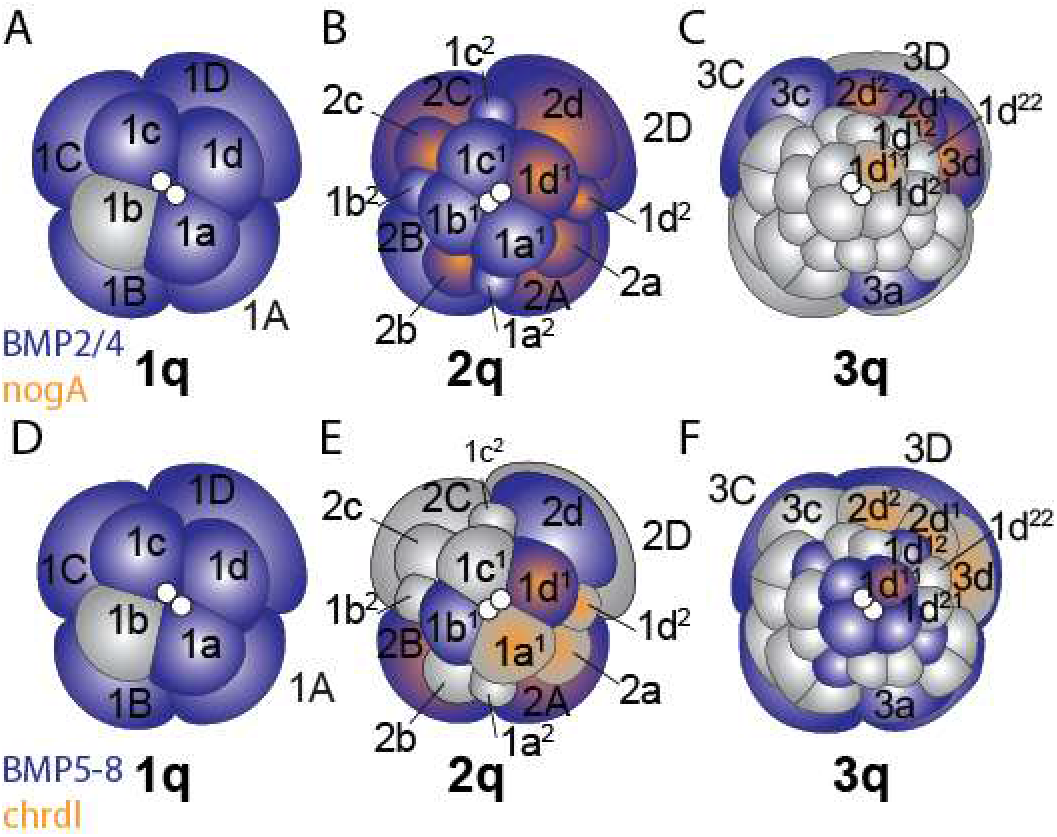
BMP ligand and inhibitor cleavage ISH patterns from Webster et al., in prep. A-C) *Ct-bmp2/4* (blue) and *Ct-nogA* (orange) at 1q (A), 2q (B), and 3q (C). D-F) *Ct-bmp5-8* (blue) and *Ct-chrdl* (orange) at 1q (D), 2q (E), and 3q (F).

Under this model, we suggest that adding exogenous BMP protein after the birth of the 2d organizer (around 3q to 4q) causes the 1a–1c micromeres to adopt a D-quadrant fate, producing an embryo with the equivalent of four 1d micromeres. Based on the fate map, this would result in a radialized brain with ectopic neural tissue, no larval eyes, and extra juvenile eye SCs—which is what we observe. At the same time, the reduced foregut may be due to only a partial loss of 2a–2c fates as 2d does not contribute to the foregut. Instead, 2d makes the majority of the trunk ectoderm as well as the VNC, which are not disrupted. We suggest this could be explained by differences in specification between the episphere and the trunk (Carrillo-Baltodano and Meyer, 2017), possibly including the role of BMP signaling. Adding BMP after the birth of 4d (until a few hours later) may result in B-quadrant micromeres adopting a C-quadrant fate, causing daughters of 1b to produce a third, ectopic, ventromedial brain lobe, larval eye, and juvenile eye SC, while 2b produces an ectopic medial foregut lobe. We were able to confirm that the third, ectopic, ventromedial eye and brain lobe are formed primarily by the 1b lineage, although contributions of 2b to the third foregut lobe were not assessed (Fig. 12).

There is support for the ability of BMP signaling to change the quadrant-specific identity of micromeres in the mollusc *Tritia* (*=Ilyanassa) obsoleta*, where a 2b lineage-specific gene *Io2bRNA* was identified (Lambert et al., 2016). After knockdown of BMP2/4 (IoDpp) with a morpholino, *Io2bRNA* was present in a subset of 2^nd^-quartet daughter cells in all four quadrants (A-D), suggesting a fate switch where daughters of 2a, 2c, and 2d adopted a 2b fate. Furthermore, after adding recombinant BMP4 protein, pSMAD1/5/8 activity appeared stronger in 1b^2^, suggesting an increased level of BMP signaling in the B quadrant (Lambert et al., 2016). Finally, in *T. obsoleta*, eye specification is based on proximity to the D-quadrant such that 1a and 1c, which are adjacent to the D quadrant, normally form eyes, but if 1b is moved closer to the D quadrant, it can form an eye (Sweet, 1998). Here 1d does not normally form eyes due to factors inherited from the polar lobe.

Our data support a role for BMP in fate switching in the episphere during early cleavage; here we either caused A- and C-quadrants to adopt a D fate around 3q, or the B-quadrant to adopt an A- or C-fate after 4q. We suggest that BMP signaling is important for establishing the quadrant identities of early blastomeres in *C. teleta*, and is a type of early organizing activity, but one that is separate from D-V axial organizing signal. Previous data suggests axial organizing comes from Activin signaling by 2d, and is responsible for the D-V axis in *C. teleta* (Lanza and Seaver 2018). We think that the blastomere trans-fating caused by addition of BMP protein seen here is occurring at a later time point (~3q) relative to the D-V organizing, and that Activin signaling may prime blastomeres to receive a BMP signal and/ or could affect the pattern of expression of the BMP pathway genes.

### BMP signaling just prior to gastrulation in *C. teleta* may pattern D-V ectodermal fates

The loss of the ventral midline and ventral shift in the expression domains of the dorsal ectodermal markers *Ct-chrdl* and *Ct-BMP5-8* that resulted from BMP treatment during the third time window (4q+12h to 4q+22h) could either be due to a shift in blastomere fates (as hypothesized for the earlier time windows) or due to alteration of an ectodermal D-V patterning system. The former hypothesis is difficult to confirm as we lack lineage tracing data for late-stage blastomeres. The ventral midline in *C. teleta* larvae largely consists of ciliated neurotroch cells that arise from descendants of 2d^12^ and 2d^2^ (Meyer and Seaver, 2010), and these neurotroch cells appear to be lost after continuous BMP treatment. In contrast, the ciliated cells around the mouth are formed by descendants of 3c and 3d (Meyer et al., 2010), and they appear unaffected by continuous BMP treatment. 2d^12^ and 2d^2^ also generate the pygidial ectoderm, telotroch, and peripheral nerves in the trunk, all of which appeared unchanged after BMP treatment. This suggests that most fates arising from blastomeres 2d^2^ and 2d^12^ are not lost after BMP treatment, although ectopic BMP could alter the fate of daughters of these blastomeres. Live imaging of daughters of 2d^2^ as well as other 2d daughter cells labeled with DiI found that the cells that will generate the neurotroch become physically separated from the cells that will generate the pygidium and telotroch early during gastrulation (Sur et al., 2020). This suggests that these presumptive neurotroch cells have received enough instruction to be different from the other 2d derivatives prior to gastrulation, possibly within our proposed time window of BMP sensitivity. We hypothesize that BMP signaling could pattern ectodermal fates along the D-V axis in the trunk prior to and possibly after gastrulation as both *Ct-BMP5-8* and *Ct-chdl* are localized to dorsal ectoderm after gastrulation. Furthermore, the loss of the ventral midline neurotroch and the ventral shift in expression of both genes after continuous BMP treatment could represent a dorsalization of the trunk ectoderm. Similar phenotypes have been reported in other annelids, which is further discussed below.

### Blocking BMP signaling in *C. teleta*

Our results highlight distinct effects of BMP signaling during different embryonic stages of development, such as the early cleavage-stage effect on quadrant-specific fates arising from the micromeres, e.g. brain and eye fates. These gain-of-function experiments did not reveal an effect of ectopic BMP4 protein on overall D-V axis formation or in blocking neural fate specification, as has been observed in some other bilaterian taxa. Previous work in *C. teleta* blocked BMP signaling using either the pharmacological agent dorsomorphin dihydrochloride or anti-sense morpholinos against SMAD1/5/8 (Joyce, 2017; Lanza and Seaver, 2018). In zebrafish, dorsomorphin has been shown to inhibit BMP type 1 receptors ALK2 (ActivinR1) and ALK3/ALK6 type 1 receptors (BMPR1), resulting in reduction of signaling by both BMP2/4 and BMP5-8 as well as Activin-A (Yu et al., 2008). Incubating cleavage-stage embryos of *C. teleta* in different concentrations of dorsomorphin produced a range of phenotypes (Joyce, 2017; Lanza and Seaver, 2018), which are described in more detail below. Overall, treatment with 5 μM dorsomorphin during cleavage stages or gastrulation to early larval development did not greatly disrupt D-V axis formation in the trunk or cause an expansion of the VNC, again reinforcing the conclusion that BMP signaling is not part of the D-V axial organizer signal or the neural inducing signal in the trunk.

Lanza and Seaver (2018) treated either 4-cell or 3q embryos for 3 h with 5 μM dorsomorphin (10 μM was tested but the resulting phenotypes were not described). They reported four separate phenotypes with a gradual reduction of features, from relatively wild-type to small disorganized balls with an episphere but no trunk. ~75% of the treated larvae had A-P and D-V axes including a brain and VNC while ~25% of the treated larvae were small, disorganized balls with anterior features including a putative radialized brain and a single ciliary band, but no trunk, VNC, or foregut. Joyce (2017) tested 5 and 10 μM dorsomorphin starting at 1q or 3q (12 h pulse) or at late cleavage or early gastrulation (continuous treatment) until stage 6. Almost 100% of animals (7 replicates) treated with 10 μM dorsomorphin at 1q arrested during cleavage and did not gastrulate while ~10-50% of animals (depending on drug lot) treated with 10 μM dorsomorphin at 3q also cleavage-arrested. Of the remaining animals, phenotypes in the nervous system, musculature, and foregut were found when treating with dorsomorphin, and many of these phenotypes were reductions in amount of tissue present.

Joyce (2017) found that animals treated with 5 μM dorsomorphin at 1q had a smaller brain (*Ct-elav1*^+^ area) with fewer 5HT^+^ neurons and a smaller, radialized foregut. In fact, ~50% of animals treated at 1q did not express *Ct-elav1* in the episphere. In contrast, animals treated with dorsomorphin at 3q did not have a significant reduction in number of 5HT^+^ neurons or foregut area, although there was a slight decrease in brain area (*Ct-elavi*^+^ area), and the foregut appeared bilaterally-symmetric in many animals. These 1q-dorsomorphin phenotypes could be consistent with a shift towards B-quadrant micromere fates and/or a loss of A/C and D-quadrant fates (less brain and foregut tissue with a lack of bilateral symmetry). However, we would also expect to see similar phenotypes with application of dorsomorphin after birth of the 3^rd^-quartet of micromeres since our first proposed time window of BMP signaling is between 3q and birth of 4d. The lack of brain and foregut phenotypes in the animals treated with dorsomorphin at 3q could be due to a delay between application of the drug and when BMP signaling is reduced. It is also unclear to what degree and how long dorsomorphin blocks BMP signaling in *C. teleta*. Finally, since neither study quantified the position of 5HT^+^ neurons in the brain nor the presence and/or localization of larval and juvenile eyes after dorsomorphin treatment, interpretation of these phenotypes relative to the blastomere-fate-switching hypothesis (proposed in this paper) are difficult.

Similar to Lanza and Seaver (2018), Joyce (2017) did not find much disruption to D-V axis formation in the trunk in animals treated with dorsomorphin at either 1q or 3q; the foregut, mesodermal bands, VNC, and neurotroch were all in the correct positions relative to one another. *Ct-elav1* expression in the trunk (i.e. VNC tissue) was reduced in 1q and 3q-treated animals in comparison to controls—including completely lacking in some animals, although when present the pattern of expression was bilaterally-symmetric with two domains of expression on either side of the ventral midline. However, the 5HT^+^ longitudinal neurites in the trunk were often quite disorganized after dorsomorphin treatment, suggesting that BMP signaling could be involved in patterning of the VNC.

Lanza and Seaver (2020a) tested the effect of SMAD1/5/8 anti-sense morpholinos (MOs) in *C. teleta*, and found a milder phenotype than with dorsomorphin: the majority of larvae had all three body axes, and 83% and 96% of larvae injected with one of two splice-blocking MOs were scored as having a D-V axis. The episphere of the SMAD1/5/8 MO-injected animals was largely normal, comprising two eyes, bilateral brain lobes, and dorsally-localized sc^ac+^ cells. Overall, most of the disruption occurred in the trunk, with a reduced neurotroch and musculature and a disruption in segmentation and formation of the ganglia of the VNC, although longitudinal connectives of the VNC were present near the ventral midline. We think the milder phenotype caused by the SMAD1/5/8 MOs could be due to a number of factors. First, while ectopic BMP4 had a dramatic effect on the amount of nuclear pSMAD1/5/8 detected (Fig. 2), the SMAD1/5/8 splice-blocking MOs did not completely block SMAD splicing (2020a). It’s possible there was still sufficient SMAD1/5/8 levels to define A/C quadrant identities. Secondly, our western blots suggest that there may be two SMAD1/5/8 transcripts produced, one of which is closer in size to homologs in other animals that retain the MH1 domain. Lanza and Seaver’s (2020) splice-blocking MOs should bind after the predicted truncation in the MH1 domain, and thus may not efficiently block the putative longer SMAD1/5/8. Overall, different experiments looking at the BMP pathway in *C. teleta* highlight the complicated nature of BMP signaling and suggest that more work is required to understand its role in development of this animal.

### BMP signaling during micromere formation in molluscs and annelids has a variable role in neural induction and D-V axis formation

The function of BMP signaling in D-V axis formation and ectodermal and neural development has been examined in ten spiralian species to date: four annelids, two brachiopods, and four gastropods (Tables 1 and 2). In the annelid *Chaetopterus pergamentaceus*, no phenotypes were detected after treatment with either 20 μM DMH1 (dorsomorphin homolog 1; inhibits the ALK2 BMP receptor) or 20 μM dorsomorphin from 4-cell to late cleavage or from late cleavage to early gastrula, making it unclear if BMP signaling was effectively blocked by either drug, and these treatments were not discussed further (Lanza and Seaver, 2020b). In two annelids and two molluscs, *C. teleta* (this study, Lanza and Seaver, 2020b, 2018), *Crepidula fornicata* (Lyons et al., 2020), *Tritia obsoleta* (Lambert et al., 2016), and *Lottia goshimai* (Tan et al., 2020), the function of BMP signaling was studied during cleavage stages, and results showed striking phenotypic similarities as well as some differences. In all four species, adding recombinant BMP4 during cleavage stages resulted in phenotypes in the episphere with very little disruption of trunk development (Table 1). All four species showed changes in the number of eyes, although the number and position of ectopic eyes varied. Interestingly, the resulting phenotypes from treating with ectopic BMP starting around or just after D-V axial organizer signaling were most similar between *C. teleta* and *Crepidula fornicata*, while *T. obsoleta* showed an almost opposite result. A loss of larval eyes was seen in *C. teleta* and *Crepidula fornicata*, whereas *T. obsoleta* showed an increase in the number of larval eyes or eyes that were closer together. Furthermore, in both *Crepidula fornicata* and *C. teleta*, ectopic BMP treatment after D-V axial organizer signaling resulted in radialization of the episphere but not of the trunk, while in *T. obsoleta*, ectopic BMP caused phenotypes in the episphere but not in the rest of the body.

**Table 1.**
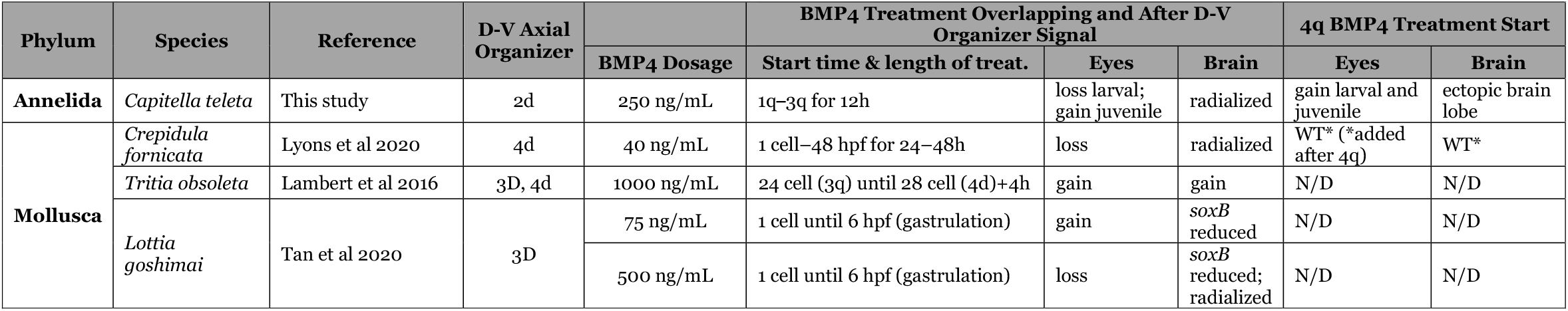
Published effects of early BMP exposure. WT: Wildtype; N/D: not determined.

**Table 2.** Published effects of BMP knockdown in spiralians. WT: Wildtype; N/D: not determined.

| Phylum | Species | Reference | Method | BMP Knockdown Phenotypes |  |  |
| --- | --- | --- | --- | --- | --- | --- |
|  |  |  |  | Neural | D-V Axial Organizer Effect | Foregut |
| **Annelida** | *Capitella teleta* | Joyce 2017 | Dorsomorphin | reduced brain & VNC | no | loss |
|  |  | Lanza and Seaver 2018 | Dorsomorphin | N/D | no | N/D |
|  |  | Lanza and Seaver 2020a | SMAD1/5/8 morpholino | wt brain, disorganized VNC | no | N/D |
|  | *Helobdella* sp (Austin) | Kuo and Weisblat, 2011 | BMP5-8 morpholino | N/D | ventralization of segmental ectoderm | N/D |
|  | *Chaetopterus pergamentaceus* | Lanza and Seaver 2020b | dorsomorphin, DMH1 | WT | WT | WT |
| **Mollusca** | *Crepidula fornicata* | Lyons et al 2020 | DMH1 | expanded brain, ectopic eyes | no | gain |
|  | *Tritia obsoleta* | Lambert et al 2016 | BMP2/4 morpholino | eyes lost | yes | loss |
|  | *Lottia goshimai* | Tan et al 2020 | Bmp2/4 morpholino | eyes lost | radialization | N/D |
|  |  |  | Chordin morpholino | ectopic eyes | radialization | N/D |
|  | *Crassostrea gigas* | Tan et al 2018 | Dorsomorphin | N/D | ventralization of gene expression patterns | N/D |
| **Brachiopoda** | *Novocrania anomala* and *Terebratalia transversa* | Martín-Durán et al 2016 | DMH1 | expansion of markers for anterior ectoderm | ventralization of ectoderm | N/D |

Although the results from *T. obsoleta* appear different than for *C. teleta* or *C. fornicata*, Lambert et al. (2016) did not test the effects of adding ectopic BMP protein later during cleavage. The increase in neural tissue and ectopic eyes seen in *T. obsoleta* treated with BMP protein from 3q until four hours after birth of 4d is similar to the formation of a third, ectopic brain lobe and third larval eye by 1b in *C. teleta* when BMP protein was added starting after birth of 4d. Furthermore, Lambert et al. (2016) reported that the phenotypes in the episphere were consistent with a loss of 1b fates after BMP soaking. The effects of ectopic BMP protein during late cleavage were also tested on *Crepidula fornicata*, but no phenotypes were seen when protein was added after birth of the 4q micromeres. It is worth noting that the phenotype of a third brain lobe and eye in *C. teleta* was not seen if BMP protein was added before birth of 4d; the radialization of brain and loss of larval eyes seemed to be a dominant phenotype. We interpret this as an inability for 1q micromeres that have been trans-fated to adopt a 1d fate by early/high BMP signaling to later change to a 1a/1c fate.

In *L. goshimai* (Tan et al., 2020), two different concentrations of ectopic BMP resulted in distinctly different phenotypes. A higher concentration of BMP protein from fertilization until the onset of gastrulation resulted in a loss of eyes and a reduction of expression of the early neural marker *soxB* in the episphere. In contrast, a lower concentration of BMP protein over the same time window resulted in ectopic eyes and a less severe reduction in *soxB* expression (Table 1). A range of BMP concentrations was also tested in *C. fornicata*, but no phenotypic differences were seen in animals treated with 2.25 ng/ml–100 ng/ml, while concentrations of 1.25 or 0.025 ng/mL had a reduction in the penetrance of phenotypes. In *C. teleta*, BMP concentrations of 250 and 500 ng/mL gave the same phenotypes, but we did see additional phenotypes in the trunk when BMP protein was changed every 12 h instead of every 24 h. Most of the studies on spiralians discussed here tested recombinant human or zebrafish BMP4 protein, although Lambert et al. (2016) did test human BMP4/BMP7 heterodimers with no reported difference in phenotypes. Taken together, this raises the question of how differing dosages of BMP protein, type of recombinant protein used, and experimental set-up may affect phenotypes reported across species, especially when the length between relevant developmental time windows (e.g. birth of the micromeres and gastrulation) may be different (e.g., hours versus days).

Blocking BMP signaling in *C. teleta* and *Crepidula fornicata* did not greatly perturb D-V axis formation in either the episphere or trunk, while blocking BMP signaling in *T. obsoleta* and *L. goshimai* caused a radialization of the resulting animals and a loss of eyes (Table 2). In *T. obsoleta*, injection of a *BMP2/4* morpholino into zygotes caused ventralization of multiple tissues and expansion of the B-quadrant-specific gene *Io2bRNA* into all four quadrants at the blastula stage, phenocopying polar-lobe ablation. This suggests that BMP signaling from the D quadrant is a crucial part of the D-V axial organizer in this animal (Lambert et al., 2016). Similarly, in *L. goshimai*, injection of *BMP2/4* or *chordin* morpholinos into oocytes resulted in radialization, suggesting that BMP signaling is part of the D-V axial organizer in this species (Tan et al., 2020). In contrast, blocking BMP signaling in *Crepidula fornicata* up until birth of the 4q micromeres using DMH1 resulted in ectopic brain tissue and eyes (Lyons et al., 2020). Injection of *SMAD1/5/8* morpholinos into *C. teleta* did not have much effect on brain or eye formation, although the connectives in the VNC were disorganized (Lanza and Seaver, 2020a). Additionally, studies in *C. teleta* have suggested that Activin signaling from the 2d micromere is part of the D-V axial organizer in this species (Lanza and Seaver, 2020a, 2018).

Overall, while the function of BMP signaling in the D-V axial organizer in these four spiralian species is different, there are also similarities in the tissues that BMP affects. We suggest that BMP signaling may affect the quadrant-specific identity of at least the 1^st^-quartet micromeres in all four of these spiralian species, as we have suggested for *C. teleta* here. Specifically, BMP signaling, possibly from the D quadrant appears to promote A/C-quadrant fates while a lack of BMP pathway activation leads to a B-quadrant fate. It is worth noting that in *C. teleta*, D-quadrant micromeres do not preferentially contribute to dorsal structures as they do in some mollusks. For example, in *C. teleta* the 1d micromere forms the dorsal-left quadrant of the episphere while 1c forms the dorsal-right quadrant, with neither micromere forming more of the dorsal tissue. The same pattern holds for the trunk—daughters of 2d generate regions of trunk ectoderm that are subdivided along the L-R and A-P axes, but not the D-V axis. These differences in contribution to dorsal tissue may explain why knocking down BMP signaling in *C. teleta* does not phenocopy a loss of the D-V axial organizer.

Other similarities in the function of BMP signaling in annelids and mollusks include the results that neural tissue in the episphere is more likely to be affected than neural tissue in the trunk, although the specific phenotype upon BMP exposure varies (Table 1). Eye formation is also affected, although losses and gains occur in different time windows and at different dosages between species. These interspecific differences in how BMP affects eyes and brain formation may be due to experimental differences as well as differences in when D-V axial organizer signaling occurs, the time windows of BMP sensitivity for individual blastomeres, and the fates generated by each blastomere.

Data from other spiralians (Tables 1 and 2) support our conclusion that BMP signaling plays an important role in brain formation, but does not appear to block neural specification in spiralians as in vertebrates and insects. Our data also support the hypothesis that neural specification occurs via two different mechanisms in the episphere and trunk of *C. teleta* (Carrillo-Baltodano and Meyer, 2017). Results reported here also align with multiple lines of evidence showing that the function of BMP signaling in neural induction is not conserved across Bilateria (Martín-Durán et al., 2018; Zhao et al., 2019). Further examination of BMP signaling in the context of spiralian D-V axial organizer and specification of blastomere quadrant-identity is warranted, especially in additional species to understand how the function of BMP has evolved across this diverse group of animals.

### BMP signaling patterns D-V fates in the ectoderm of multiple spiralians

In spiralians, a later role of BMP signaling after gastrulation has also been studied. Within Pleistoannelida (Errantia + Sedentaria) (Struck, 2011), three taxa have been examined: *C. teleta*, *Helobdella sp*. Austin, and *Platynereis dumerilii*. Data from *P. dumerilii* also suggest a function of BMP signaling in patterning ectodermal and neural fates along the D-V axis in the trunk. Denes et al (2007) found that exogenous BMP4 added after gastrulation (24hpf, trochophore stage) resulted in a downregulation of some ventral markers such as *Pdu-nk2.2* and the midline marker *Pdu-sim* (*single-minded*) and an expansion of dorsal markers such as *Pdu-pax3/7*, although the domains of the ventrolaterally-expressed genes *Pdu-elav* and *Pdu-pax6* were not affected. In *Helobdella sp*. Austin, gain- and loss-of-function experiments in the teloblast cells demonstrated that BMP signaling is important for D-V patterning of the ectoderm within the midbody and caudal segments; however, no effects on neural induction or the overall D-V axis including the mesoderm were reported (Kuo et al., 2012; Kuo and Weisblat, 2011). Anti-sense *Hau-BMP5-8* morpholinos caused a dorsal-to-ventral shift in segmental ectodermal fates; p bandlet cells adopted the more ventral o bandlet fate. In contrast, ectopic expression of *Hau-BMP5-8* mRNA in n bandlet cells caused the neighboring o bandlet cells to adopt a more dorsal p bandlet fate; o bandlet cells upregulated *Hau-six1/2a* and down-regulated *Hau-pax6a*. Interestingly, *Hau-pax6a* expression in the n bandlet cells themselves, which largely contribute to the VNC in leech, was not affected by ectopic BMP signaling. These findings are similar to our results in *C. teleta*, where BMP4 treatment just prior to gastrulation caused a ventral expansion of *Ct-chrdl* expression in the dorsal ectoderm and a loss of the ventral midline neurotroch but not the VNC. Taken together, these data suggest a later function of BMP signaling in patterning dorsal ectodermal fates in the segmented body (trunk) of annelids, although the amount of neuroectodermal tissue appears to be largely unaffected by perturbations of BMP across all three annelids tested.

A function of BMP signaling in patterning D-V fates in the ectoderm has also been shown in other spiralians. In the mollusc *Crassotrea gigas* (Tan et al., 2018), dorsomorphin treatment (0.32 μM from 3–5 hpf) caused ventralization; expression of ventral genes was expanded while dorsal gene expression was reduced. In the brachiopods *Novocrania anomala* and *Terebratalia transversa* (Martín-Durán et al., 2016), DMH1 caused an expansion of some ventral ectodermal genes, as well as an expansion of anterior genes and reduction of posterior ectodermal genes. Finally, during regeneration in the flatworm *Dugesia japonica* (Orii and Watanabe, 2007), *DjBMP* RNAi produced an ectopic ventral side. Together, these data suggest a possible function for BMP signaling in patterning the ectoderm across spiralians.

### Conclusions

Adding exogenous BMP protein to early-cleaving *C. teleta* embryos had a striking effect on formation of the brain, eyes, foregut, and ventral midline in a time-dependent manner. However, adding BMP did not block neural specification of the brain or VNC or block formation of the D-V axis at any of the treatment time windows. Strikingly, there was little to no reduction in neural tissue in the VNC after these continuous BMP treatments. Our results, in conjunction with data from other spiralians, suggest that BMP signaling was not ancestrally involved in delimiting neural tissue in the last common ancestor of spiralians. However, BMP signaling may have played an organizing role in establishing blastomere identity in the last common ancestor of annelids and mollusks, although this function appears to be separate from and possibly later than the D-V axial organizer in some spiralian species. Furthermore, BMP signaling may have functioned to pattern ectodermal fates along the D-V axis in the trunk of the last common ancestor of annelids. Ultimately, studies on a wider range of spiralian taxa are needed to determine how variations in the role of BMP signaling in blocking neural induction and establishing the D-V axis have evolved. These comparisons will give us insight into the evolutionary origins of centralized nervous systems and body plans.

## Acknowledgements

The authors thank R. Bellin at the College of the Holy Cross for access to the confocal microscope and the two anonymous reviewers for their in-depth, helpful comments.

